# Microwell-Based Flow Culture Increases Viability and Restores Drug Response in Prostate Cancer Spheroids

**DOI:** 10.1101/2022.06.28.498007

**Authors:** Marie C. Payne, Sum Yat Ho, Takao Hashimoto, Sara Imboden, Johnny A. Diaz, Brandon S. Lee, Melissa J. Rupert, Nathan Y. Cai, Andrew S. Goldstein, Neil Y.C. Lin

**Affiliations:** Department of Mechanical & Aerospace Engineering, University of California, Los Angeles, Los Angeles, CA 90095-1597; Department of Biochemistry, University of California, Los Angeles Los, Angeles, CA 90095-1597; Departments of Molecular, Cell & Developmental Biology and Urology, University of California, Los Angeles Los, Angeles, CA 90095-1597; Department of Bioengineering, University of California, Los Angeles Los, Angeles, CA 90095-1597; Institute for Quantitative and Computational Biosciences, University of California, Los Angeles Los, Angeles, CA 90095-1597

**Keywords:** Spheroid, Millifluidics, Prostate Cancer, Flow Culture, Imaging, Tissue Engineering

## Abstract

3D cancer spheroids represent a highly promising model for study of cancer progression and therapeutic development. Wide-scale adoption of cancer spheroids, however, remains a challenge due to the lack of control over hypoxic gradients that may cloud the assessment of cell morphology and drug response. Here, we present a Microwell Flow Device (MFD) that generates in-well laminar flow around 3D tissues via repetitive tissue sedimentation. Using a prostate cancer cell line, we demonstrate the spheroids in the MFD exhibit improved cell growth, reduced necrotic core formation, enhanced structural integrity, and down-regulated expression of cell stress genes. The flow-cultured spheroids also exhibit an improved sensitivity to chemotherapy with greater transcriptional response. These results demonstrate how fluidic stimuli reveal the cellular phenotype previously masked by severe necrosis. Our platform advances 3D cellular models and enables study into hypoxia modulation, cancer metabolism, and drug screening within pathophysiological conditions.

## 1 Introduction

3D cancer spheroids are valuable *in vitro* models that have been extensively used in both fundamental research and industrial settings, ranging from precision medicine and drug development to cellular therapies [1, 2]. For example, analysis of cellular growth and migration in *in vitro* cancer spheroids has revealed mechanistic pathways underlying invasive tumor development and metastasis [3, 4]. Studies of multicellular cancer spheroids have also shown that cellular cross-talk plays a role in tumor growth and tumor-mediated immune suppression [5–8]. More importantly, reconstructing the native 3D architecture has led to greater therapeutic insight into radio- and chemo-sensitivity generated by limited drug penetration and reduced cellular proliferation [9, 10]. In these studies, the 3D spheroids better recapitulate the cell-cell and cell-matrix interactions within tumors compared to traditional 2D monolayer culture [11–13]. These interactions closely mimic the complex physical cues found in native tissues and, in turn, lead to *in vivo*-like gene expression profiles and drug responses [14–18].

Despite such an improved reconstruction of the tumor microenvironment, there is still a substantial discrepancy between 3D culture and primary tumors. A fundamental limitation to 3D spheroids is the development of steep oxygen and metabolite gradients within spheroid cores, leading to necrosis [19–21]. Solute transport within avascular tissues typically relies on passive diffusion, which restricts nutrient exchange to cells beyond the oxygen diffusion limit of ~ 100-200 *μ*m [22, 23]. While a well-controlled hypoxic condition in *in vitro* models can be useful to study limited tumor growth, hypoxia-induced cell invasion, drug resistance development, and cellular adaptations to oxidative stress [24–27], conventional 3D spheroid systems usually exhibit overly severe and physiologically irrelevant levels of necrosis, causing a strongly biased understanding of therapeutic efficacy for three major reasons [28]. First, the sizeable necrosis in large spheroids impacts many essential cellular processes such as the penetration, binding, and bioactivity of therapeutic drugs and drug candidates [29, 30]. Second, most *in vivo* hypoxic conditions are transient, whereas the hypoxic condition in spheroids continuously worsens as the sample grows [19]. Lastly, variations in cell packing and necrotic core size can dominate over cell-specific responses and challenge high-throughput screening accuracy [31, 32]. Identifying solutions for precise control over hypoxic gradients and necrotic formation would greatly improve physiological and clinical relevance in tumor spheroids.

To date, flow culture has emerged as one of the most popular methods for addressing hypoxia-induced necrosis. Microfluidic devices, bioreactors, and spinner flasks, have all proven to promote intra-tissue transport of oxygen, nutrients, and metabolic wastes [33–35]. In various cancer tissue models, the applied flow has been shown to preserve microenvironment heterogeneity, cellular viability, and drug responsiveness [36, 37]. In addition, the fluidic mechanical stimulus has been implicated in enhancing aggressive cancer phenotypes and chemoresistance [38–41]. Fluidic devices have become commonplace in tissue engineering methodology, however more innovation is required for increased performance, throughput, and adoption. Bioreactors and spinner flasks are limited in that they usually demand large quantities of media, generate non-uniform shear stress, and prohibit culture of independent replicates; furthermore, samples are not easily accessible for real time monitoring. Microfluidic systems, on the other hand, require cumbersome fabrication and operation procedures that hinder high-throughput applications [42,43]. Therefore, there is a technological gap that must be resolved for next-generation 3D tissue culture.

Here, we address challenges of hypoxia-induced necrosis in 3D prostate cancer (PCa) spheroids by developing a fluidic system, the Microwell Flow Device (MFD). Compared to macro-scale devices (i.e., orbital shakers and spinner flasks) the MFD generates laminar flow around independent replicates within millimeter scale wells. Compared to microfluidic devices, our millifluidic system offers simple fabrication procedures and can be easily scaled up to accommodate molecular screening. The MFD operates by taking advantage of the natural density differences (~10%) between biological tissue and surrounding media – by repeating a 180-degree flipping motion, the spheroid will be re-suspended within its respective well and allowed to freely sediment to the bottom, thereby generating an external flow field without the use of tubing or pumps. To demonstrate the utility of the MFD culture platform, we generated large (4k cells/well) spheroid models from the Lymph Node Carcinoma of the Prostate (LNCaP) cell line. Large spheroids achieve better physiological relevance in terms of growth, cell function, and drug responses, but have been underutilized in *in vitro* studies due to challenges in viability and long-term growth [44,45]. We investigate that LNCaP spheroids in the MFD exhibit reduced necrosis and maintain cellular structural integrity throughout the spheroid. Such a dynamic growth environment can prevent the development of toxic, irreversible oxygen gradients which may mask important cell phenotypes [11].

## 2 Results

### Design of a Microwell Flow Device for Individual, Laminar Flow Culture of Spheroids

To facilitate solute transport in the LNCaP tissues, we designed the MFD to provide uniform shear stress based on spheroid sedimentation. The system is designed to work with commercial well plates and is composed of a custom lid and clamp that is periodically rotated by a single stepper motor 180 degrees (**Figures 1a** and S1). The custom lid is formed by a rigid outer shell, rubber padding, and a 150*μ*m-thick Polydimethylsiloxane (PDMS) membrane (Figure 1b). PDMS is a silicon-based material that is bio-compatible and simple to fabricate [46] (see Figure S2 for fabrication process). Through its porous structure, the PDMS facilitates this flipping motion by simultaneously retaining liquid and allowing gas exchange to the media. The rate of gas exchange is thickness dependent and is comparable to the rate of exchange in an open-diffusion plastic lid (Figures 1c and S3).

**Figure 1:**
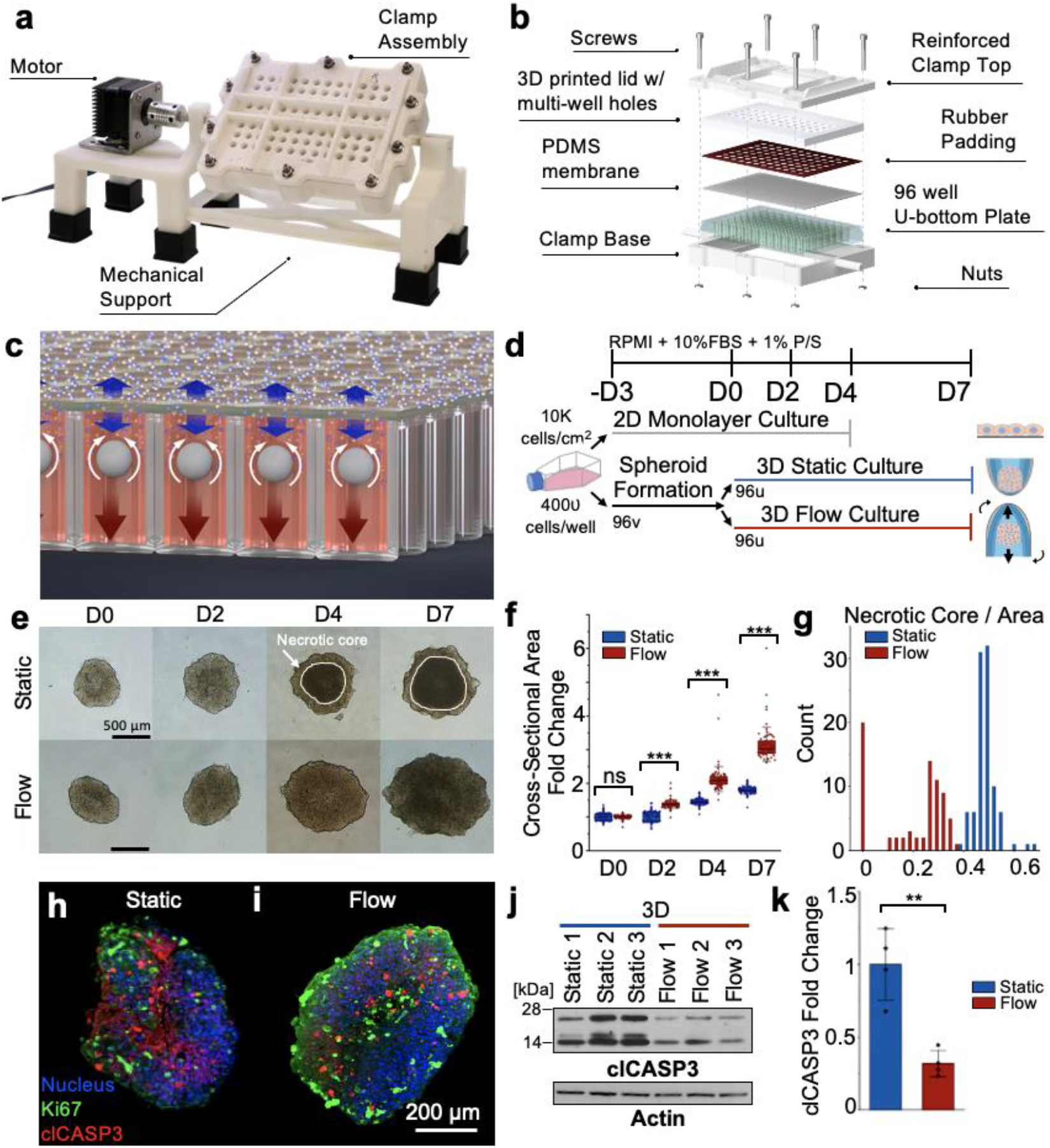
MFD fabrication and PCa spheroid growth. **a** Major components of the MFD include a stepper motor, clamp assembly, and mechanical support. **b** Schematic of the custom clamp design, featuring rubber padding and a 150 *μ*m PDMS membrane to facilitate gas exchange while retaining media. The lid and plate are secured by a 3D printed clamp with 10 exterior screws. **c** Schematic of spheroid motion and gas exchange through the PDMS membrane in individual wells. **d** Timeline of spheroid and 2D monolayer culture. **e** Brightfield microscopy images of flow and static samples. The white outline delineates the formation of the necrotic core in static that is absent in the flow sample. **f** Growth quantification of spheroids from brightfield images (static min n = 85; flow min n = 73 biological replicates (BRs)). **g** Measurements of the necrotic core to cross-sectional area ratio on D7 (static n = 95; flow = 73 BRs). **h-i** Immunofluorescent imaging of cellular proliferation (Ki67) and apoptosis (clCASP3) in D7 spheroids. **j** Western Blot analysis of clCASP3 with Actin as loading control. **k** Quantification of relative clCASP3 protein expression (static and flow n = 4 BRs). (ns = non-significant, ** p<0.01, *** p<0.001, two-tailed Student’s T-Test.)

Additionally, the MFD addresses limitations of conventional fluidic systems. Compared to a standard 125ml spinner flask or bioreactor, each well in a 96-well plate requires a maximum of 350 *μ*l of media, which is approximately a 3.5-fold reduction in total media consumption. The elimination of mechanical stirrers and tubing resolves unwanted fluidic shear stress gradients, decreases damaging cellular collisions, and reduces labor-intensive culture maintenance [47–49]. Moreover, by seeding each spheroid in its own well, the samples may be kept biologically independent, providing enhanced statistics and throughput for assays. Using computational fluid dynamics and Stokesian analysis, we estimate a terminal superficial shear stress of ~0.16 dynes/cm^2^ for a settling spheroid with radius 500*μ*m (Figure S4), which is consistent with physiological values found in tissue interstitial flow and lymph flow [50] and has been used for modeling cancer cell invasion [51].

To test how our flow culture device influences the phenotype of 3D human cancer models, we generated spheroids using a metastatic PCa cell line (LNCaP). In brief, the spheroid samples were first generated by seeding 4k cells per well in v-bottom plates to allow the cells to aggregate. 3 days post-seeding (Day 0), the samples were split into MFD (hereafter referred to as “flow” and 3DF for 3D Flow) and static (3DS for 3D Static) conditions for 7 days (Figure 1d). On Day 0, we first observed that flow enhanced the perfusion of molecules, indicated by the increased fluorescent intensity of a neutral lipid binding dye (Bodipy) that stains lipid droplets, within 1 hour of being placed in flow or static culture (Figure S5). Using transmitted-light microscopy, we found that the 3DF and 3DS samples demonstrate substantially different growth and necrotic core formation (Figure 1e). Measurement of the cross-sectional area validates the significant size differences between the two conditions (Figure 1f). By Day 7, the 3DF spheroids had an average fold increase of 1.5 compared to 3DS, demonstrating high consistency across 4 independent runs (Figure S6). Upon comparison with traditional orbital shaking of 96-well plates, we found that our observed spheroid growth enhancement cannot be achieved by orbital shaking, reinforcing the advantage of sedimentation-induced flow in the MFD (Figure S7). In addition, 3DS spheroids develop a necrotic core that occupies an average of 50% of the total cross-sectional area on Day 7, which was reduced by ~30% in the 3DF sample (Figure 1g). Immunostaining of Ki67, a marker of cell proliferation, and clCASP3, a marker of cell apoptosis, displays a strong accumulation of clCASP3 in the static spheroid cores that is largely reduced in the flow samples (Figures 1h-i, Figure S8). The 3DF spheroids exhibited an increased outer proliferative zone, consistent with previous spheroid studies [52,53]. Furthermore, Western Blot analysis confirms the immunofluorescent results of clCASP3 (Figures 1 j-k, Figure S9); additional protein expression for hypoxia-related markers was performed, showing no statistically significant difference (Figure S9).

We then investigated the importance of continual flow culture by testing the re-emergence of the necrotic core upon flow cessation. Both flow and static spheroids were transferred into static wells on D7 and stained with NucGreen and NucBlue for dead and total cell imaging, respectively (**Figures 2a**, S10, and Video S1). After 12 hours, the flow-to-static spheroids began to show an increased NucGreen intensity throughout the sample core. This delayed onset of acute hypoxia was validated by quantifying the dead cell population (Figure 2b). Within 24 hours after being placed in static culture, the viable phenotype was effectively diminished in the flow-static samples.

**Figure 2:**
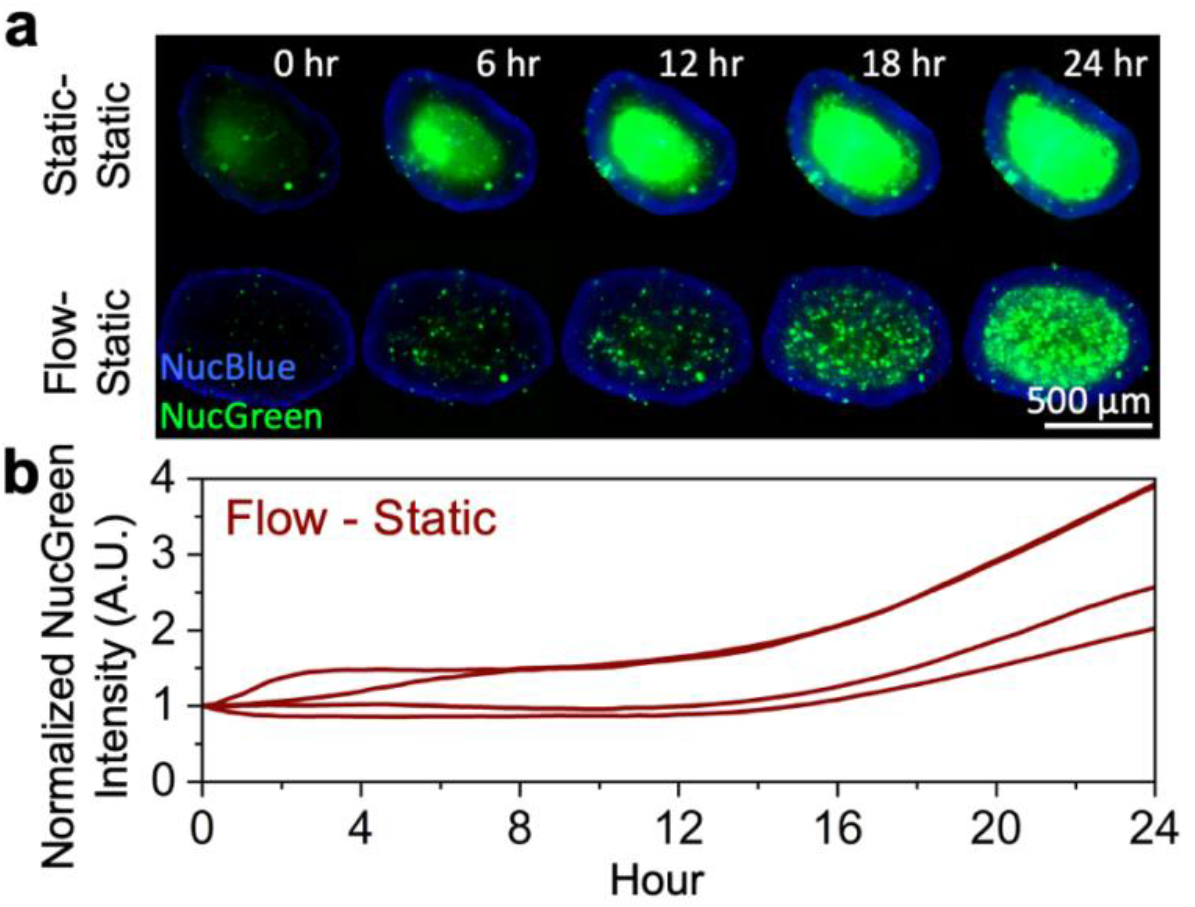
24hr onset of acute hypoxia after stopping flow. **a** Timelapse of dead (NucGreen) and nuclear (NucBlue) staining for D7 flow and static samples placed in static culture for 24 hours. **b** Quantification of dead cell population in Flow-Static spheroids by measurement of average NucGreen intensity after background subtraction (Flow-Static n = 4 BRs).

### Flow culture preserves cellular structure and behaviors in LNCaP spheroid cores

To further understand how the applied flow influences the spheroid structure and cellular phenotype, we performed immunostaining of spheroids in both flow and static culture (**Figures 3a-e**). We found that the cells in 3DF samples exhibit larger nuclei with greater cell-cell separation compared to the 3DS samples (Figures 3a, d-e). Furthermore, we found that in 3DS samples, only the peripheral cell layers exhibit well-established intercellular junctions, indicated by locally enriched E-cadherin (E-cad) expression (Figures 3b-c). In contrast, we observed well-established E-cad junctions in 3DF samples across the entire cross-section. Such a finding suggests that the flow culture uniformly preserves the E-cad-mediated cell-cell interaction, which regulates many cancer-related biological processes such as cell invasion and transdifferentiation [54]. In addition, both 3DS and 3DF samples display lipid droplets (Bodipy) and express FASN (fatty acid synthase) and vimentin (Figure S11), akin to 2D culture (Figure S12). Androgen receptor (AR) and its target prostate-specific antigen (PSA) are also expressed in both 3D samples without significant difference (Figure S11).

**Figure 3:**
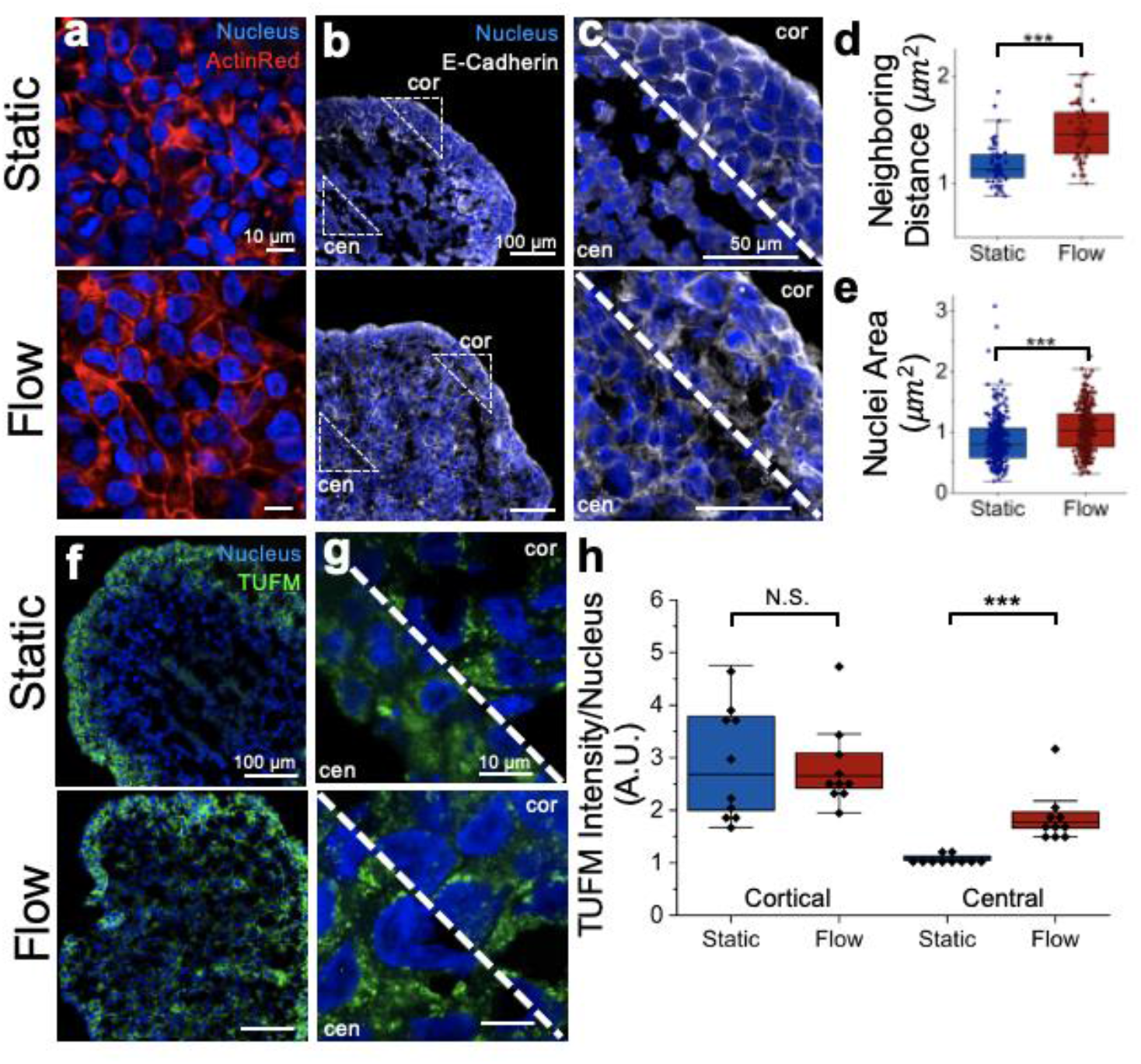
Immunofluorescent image quantification of 3D LNCaP cultures. **a** Sectioned LNCaP spheroids immunostained for Actin (ActinRed) and DNA (DAPI). **b-c** Immunostained LNCaP sections of E-cadherin (E-cad) illustrate cell boundaries in cortical (cor) and central (cen) regions. **d-e** Quantification of distance between neighboring nuclei (flow n = 50; static n = 50 BRs) and nuclear area (flow n = 275; static n = 268 BRs). **f-g** Immunostaining of TUFM reveals a breakdown in mitochondrial distribution in 3DS cen regions compared to the viable 3DS cor regions. 3DF cen and cor TUFM intensity does not display a visible morphology contrast. **h** Quantification of TUFM intensity per nucleus density (n=5 technical replicates (TRs) per BR; n=2 BRs for each of flow and static). ns = non-significant, *** p<0.001, two-tailed Student’s T-Test.

We next characterized the mitochondrial morphology and distribution by immunostaining of TUFM (Tu Translation Elongation Factor, Mitochondrial), a marker of the mitochondrial inner membrane. TUFM expression has been shown to be positively correlated with E-cad expression and plays an important role in mitochondrial function and metastatic development [55]. Between the 3D spheroid cultures, we observed a reduction of TUFM signal in the central area of the 3DS spheroids, whereas 3DF samples exhibited a uniform intensity distribution (Figure 3f). Additionally, the mitochondrial architecture appears spherical or fragmented, characteristic of proliferating cells, across 3DF and in the 3DS cortical region, unlike the 3DS central region (Figure 3g). Quantification of the TUFM intensity per nucleus validates the cortical mitochondrial similarity versus the central mitochondria loss in the 3DS samples (Figure 3h).

### Flow culture restores hypoxia-induced transcriptional regulation in LNCaP spheroids

To understand the transcriptional response of spheroids under flow, we analyzed the RNA expression level of 45 genes comprising markers of PCa, metabolism, cell viability, and hypoxia (Table S1). By analyzing D7 spheroids, we found that the three culture conditions (i.e., 2DS, 3DS, and 3DF) formed distinct clusters, indicating both the model dimensionality (i.e., 2D vs. 3D) and physiological stimulus (i.e., flow vs. static) impose individual effects on the LNCaP cell behavior (**Figure 4a**). To highlight the differential gene expression profile induced by the applied flow in the 3D conditions, we generated a volcano plot by normalizing 3DF to 3DS (Figure 4b). Our analysis showed that the flow significantly downregulates hypoxia markers *Ca9* and *Cxcr4* by more than 50%. We also observed upregulated expression of *Slc1a5,* a glutamine transporter, and *Ndufa8,* a mitochondrial complex-I subunit, which are involved in glutamine metabolism and oxidative phosphorylation [56,57]. Moreover, we found an inverse differential expression between *Eno2,* a neuroendocrine marker, and *Folh1,* the prostate-specific membrane antigen gene, which indicates a reduced neuroendocrine-like phenotype in 3DF vs. 3DS [58]. Bar charts of the differential gene expression normalized to the 2D control further illustrate how the gene expression under flow largely recovers back to the baseline expression levels seen in 2D LNCaP models (Figure 4c). These findings collectively suggest that the applied flow mitigates the hypoxia condition and in turn impacts the metabolic and neuroendocrine-like phenotypes of LNCaP cells.

**Figure 4:**
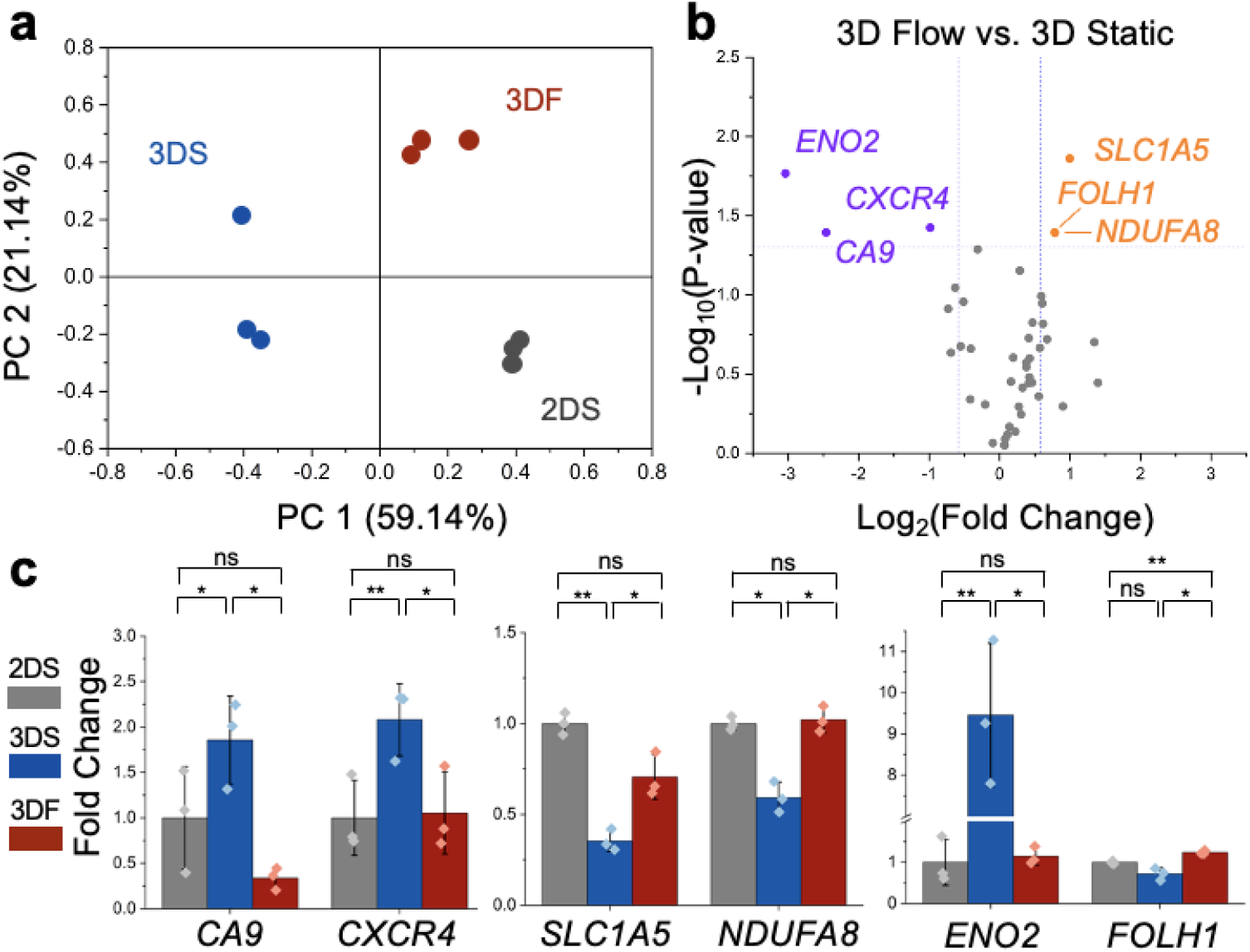
Gene expression analysis of LNCaP culture conditions. **a** Principal Component Analysis (PCA) plot of 3D Flow (3DF), 3D Static (3DS), and 2D Static (2DS) culture conditions shows clustering based on normalized gene counts for 45 tested genes (OriginPro). Each data point represents one BR. **B** Volcano plot of upregulated (orange) and downregulated (purple) genes in 3DF condition normalized to 3DS. Significance was determined at p ≤ 0.05 and fold change ≥ 1.5. **c** Gene expression bar charts of six differentially expressed genes normalized to 2DS. Data points represent one BR. ns = non-significant, * p<0.05, **p<0.01, two-tailed Student’s T-Test.

### Flow-cultured spheroids enable precise modeling of cellular responses to docetaxel

The improved phenotype of LNCaP cells in our 3D flow culture allows us to characterize the drug response that is masked by pronounced cell death. Here, we perform a dosage response assay using docetaxel, which is routinely used for treating advanced stages of PCa either alone or in combination with other drugs [59,60]. We tested four representative concentrations (0, 5, 10 and 20nM) by administering docetaxel to 3DS and 3DF samples on D5 for 48 hours (Figure S13). To visualize dose dependence in the 3D samples, we performed dead and total cell staining (**Figure 5a**) for the dead cell ratio quantification. Even without the drug applied, the static sample exhibits pronounced cell death, masking the cell toxicity arising from the docetaxel treatment. In contrast, the flow culture exhibited a low dead cell baseline in the untreated sample, similar to the 2D control, allowing for improved characterization of the dosage response (Figure S14). Such a finding is confirmed by quantifying the ratio of total to dead cells between 3DS and 3DF (Figure 5b). Further quantification of the spatial distribution of dead cells confirms most cell death occurs within the necrotic core of the 3DS samples, whereas 3DF samples exhibit a more uniform dead cell expression (Figure 5c). Our cell viability analysis illustrates the importance of intra-tissue solute transport for uncovering potential confounders in toxicity assays.

**Figure 5:**
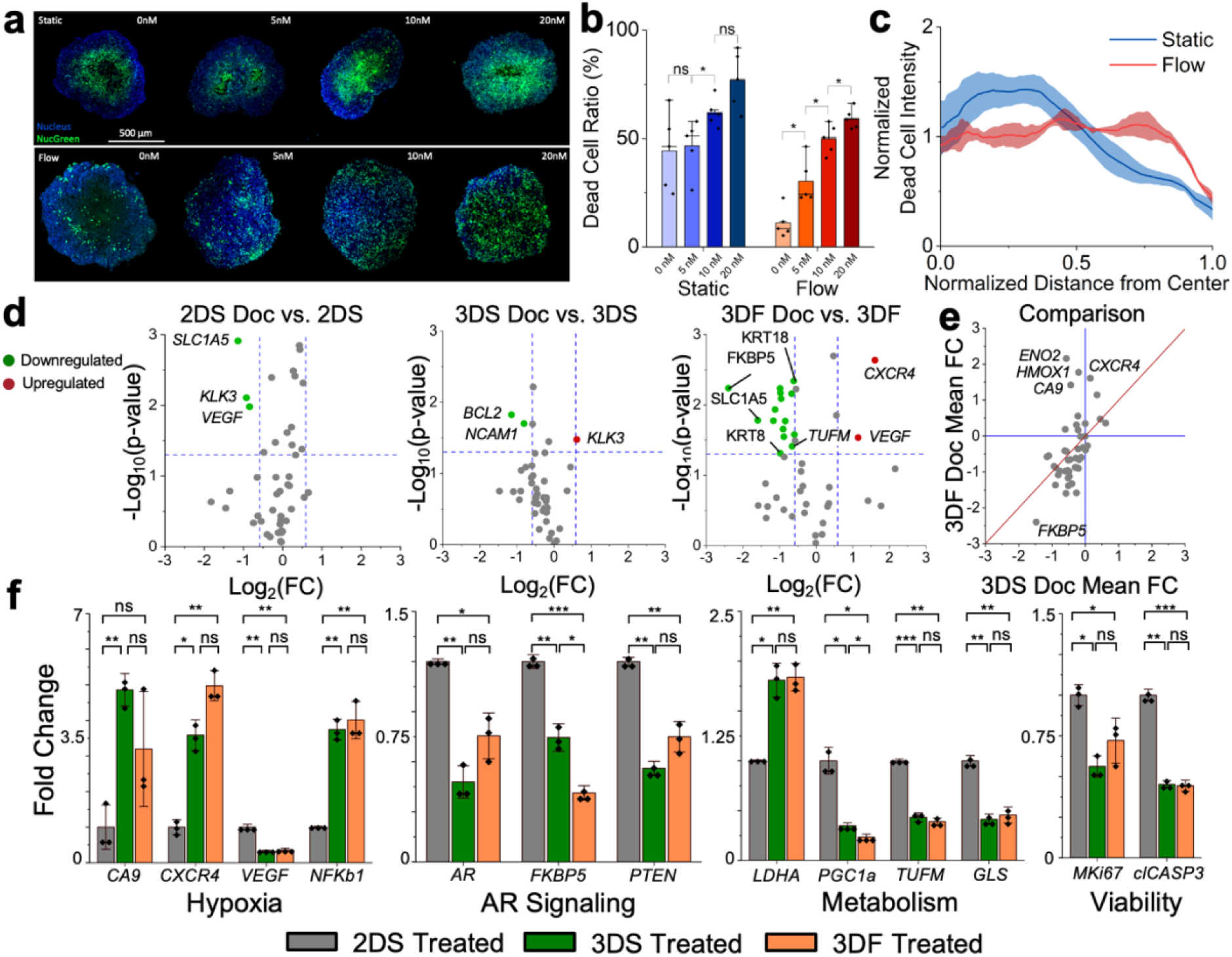
Differential gene expression of docetaxel in static vs. flow cultured spheroids. **a** Dose response of static and flow spheroids for 0, 5, 10 and 20nM concentrations of docetaxel (**a-c** static and flow n=5 BRs). **b** Measurement of the ratio of dead cells to the entire spheroid area. **c** Dead cell distribution by normalized azimuthal integration of NucGreen intensity from the center to the perimeter of the spheroid. **d** Volcano plots of 2DS, 3DS and 3DF differential gene expression for treated samples normalized to corresponding untreated samples. Significance was determined at p ≤ 0.05 and fold change ≥ 1.5. Each assay was performed in triplicate. **e** Comparison of mean fold change between 3DS and 3DF treated vs. untreated samples. Genes in the second quadrant indicate upregulation in flow samples and downregulation in static samples post treatment. **f** Bar charts of differential gene expression of treated samples for markers encompassing hypoxia, AR signaling, metabolism, and cell viability. Each data point represents one BR. ns = non-significant, * p<0.05, ** p<0.01, *** p<0.001, two-tailed Student’s T-Test.

To further understand how the culture condition impacts the cellular response to drug treatment, we visualized gene expression changes in 10nM docetaxel treated vs. untreated samples using volcano plots (Figure 5d). Compared to the 2DS and 3DS samples, the 3DF spheroids reveal more differentially expressed genes. To identify the effect of flow on docetaxel-induced transcriptional response, we generated a comparison of the gene expression fold change between treated and untreated conditions in flow and static (Figure 5e). This analysis shows that the docetaxel treatment upregulates essential hypoxia (i.e., *CA9, CXCR4,* and *HMOX1*) and neuroendocrine (i.e., *ENO2)* markers in the flow sample. This result, however, is masked in the static sample, since these hypoxia-related markers readily exhibit a high baseline expression level in untreated spheroids. Furthermore, the expression level of genes related to hypoxia, AR signaling, metabolism, and cellular viability becomes non-significant between both 3D drug-treated samples (Figure 5f). Together, our results suggest the interpretations of cell response to drug treatment heavily relies on physiologically relevant growth conditions.

## Discussion & Conclusions

3D tissues better recapitulate *in-vivo* tissue growth and therapeutic response compared to 2D culture however there are technical hurdles that prevent widespread adoption. Primarily, the insufficient solute transport causes overwhelming hypoxia-induced necrosis in the core of spheroids, generating steep metabolite and oxygen gradients across the sample, clouding the assessment of cell phenotype and drug response [61,62]. In this study, we report a Microwell Flow Device (MFD), a scalable system that generates physiologically relevant fluidic stimuli that drives nutrients toward the tissue core, which in turn maintains central tissue structure and cellular behaviors. Our system generates laminar flow and individual replicates, two qualities that have been difficult to achieve simultaneously in 3D flow culture. The MFD can be tailored to any plate dimension and, in future work, may be scaled up to accommodate mid-throughput (*>*1000 sample) assays. These properties serve to improve tissue quality and uniformity as well as compatibility with high throughput screening platforms, thereby encouraging use of 3D tissue models for both basic research and therapeutic development.

As demonstrated in LNCaP spheroids, flow-cultured samples can grow larger for longer periods of time without excessive central necrosis. On the cellular level, we observed reduced necrosis throughout the spheroid and increased cellular proliferation near the sample periphery. Immunofluorescence microscopy reveals restoration of cell packing, intercellular adhesion, and mitochondrial morphology in flow samples. On a transcriptional level, gene expression measurement shows that the applied flow reverts the expression of hypoxia, metabolic, and neuroendocrine-related markers that were altered in 3D static culture. Lastly, the MFD spheroids showed increased sensitivity to docetaxel compared to the 3DS samples, with a uniform cell death distribution and higher differential transcriptional response. This finding is consistent with previous studies which suggest that removing hypoxia-induced cellular drug resistance could increase the cellular response to chemotherapy [63–65], and which show flow culture-derived spheroids can reduce hyper-sensitivity to therapeutics commonly seen in 2D cultures [66].

Our culture platform further provides a foundation for mechanistically investigating longstanding questions in PCa. First, cancer metabolic activity plays a pivotal role in tumor growth, metastasis, and drug response, but can be strongly affected by hypoxia. [25, 67, 64]. Our MFD effectively restores the cellular behavior in 3D spheroids, providing an improved platform for metabolic assays. Furthermore, by increasing the complexity of the spheroid microenvironment, our system can enable studies that provide insights into intercellular signaling and how it dictates tumor remodeling. For example, multicellular and hydrogel-embedded models could offer added control over the tumor microenvironment [68]. Similarly, vascularized spheroids, organoids, and patient-derived explants [69], can be directly implemented in our platform to better recapitulate the native tissue responses. By growing independent replicates, our device provides a scalable platform for compound screening, accelerating developments of combinatorial therapy and precision medicine.

## 3 Methods

### MFD Manufacturing

All components of the MFD were 3D printed (Formlabs Form 3). The custom 96 well plate lid consisted of a PDMS membrane and a laser-cut silicone rubber sheet (McMaster-Carr, 9010K11) adhered to the lid using silicone sealant (Loctite, 908570). For the PDMS membrane, 10g of Sylgard 184 (Dow Corning, 11-3184-01 C) was mixed with a 10:1 monomer:catalyst ratio, spin coated to a thickness of 150*μ*m, and cured at 150°C for 35 minutes. Prior to use, all lid components were sterilized by soaking in 70% ethanol for a minimum of 10 minutes before use and air dried. Device rotation was controlled by a NEMA 17 stepper motor connected to an A4988 stepper motor driver and Arduino UNO microcontroller, programmed to flip 180° every 10 seconds.

### Cell Culture

Human prostate adenocarcinoma-derived LNCaPs were cultured according to the ATCC thawing, propagating, and cryopreserving protocol [70,71]. The culture media for LNCaP comprised of RPMI 1640 (Gibco), 10% fetal bovine serum (Gibco), and 1% Penicillin/ Streptomycin (Gibco). LNCaP culture was incubated at 37°C, 5% CO2, and 90% relative humidity. Media change was performed every 24 to 48 hours. Subculture of LNCaPs was performed at ~ 80% confluency, in which the cells were washed with 1X PBS -/- twice and incubated with 0.5% Trypsin-EDTA at 37°C for cell detachment. The dissociated cells were then centrifuged at 250g for 3 minutes and re-suspended in warmed culture media. In 2D culture, the seeding density was 10,000 cells/cm2 on poly-l-sine (Sigma-Aldrich) coated surface. For 3D culture, LNCaPs were seeded into Greiner Bio-One™ CellStar™ 96-Well, Cell Repellent-Treated, V-Shaped Bottom Microplate at 4K cells/well. For better aggregation of LNCaP, the 96-well plate is centrifuged at 300g for 10 minutes. Before introduction of the flow or drug conditions, the spheroids were allowed to grow for 72 hours post seeding. For experiments, LNCaP spheroids were transferred into Corning 96 Well Clear Ultra Low Attachment Microplates. To monitor the growth of monolayer and spheroid culture, phase contrast images were taken right after media change on an Olympus CKX41 at 4x magnification.

### Spheroid Sectioning & Immunofluorescent Imaging

Spheroids for cryosectioning were fixed in 4% Paraformaldehyde (PFA) in Phosphate Buffered Saline (PBS +/+) for 20 minutes on ice and then washed three times in PBS +/+. Spheroids were then placed in 30% sucrose in PBS +/+ for 1-3 hours on ice until completely submersed. The spheroids were embedded and frozen in Tissue-Tek Optimal Cutting Temperature (O.C.T., Sakura) compound, cryosectioned at 12*μ*m thickness, and collected onto Superfrost Plus slides (Fisher Scientific). The LNCaP samples were first blocked using a mixture of 2% donkey serum (Sigma-Aldrich, D966310ML) and 0.5% Triton X-100 (Sigma-Aldrich, T8787-50ML) in PBS +/+ for 30 minutes (2D) or 60 minutes (whole & cryosectioned spheroids). After blocking, the slides were washed with PBS twice, and then incubated with the primary staining solution (0.5% BSA, 0.25% Triton X-100, and the primary antibody (SI Table 2)). The samples were left in the staining solution for 30 minutes (2D), overnight (cryosectioned slides), or for 24 hours (whole spheroids), followed by two washes with 1X PBS. Afterwards, the secondary staining solution (with NucBlue and the secondary antibody (SI Table 2)) was added for 30 minutes (2D), 60 minutes (cryosectioned spheroids), or 24 hours (whole spheroids). 2D and whole mount spheroids were washed twice with PBS and stored in 0.1% Tween 20 (Sigma-Aldrich, P9416-50ML) at 4°C. Sectioned slides were mounted with cover slips using Prolong Diamond antifade mountant (Thermo Fisher Scientific, P36970) and stored at 4°C.

### Gene Expression Measurements

To prepare 3D LNCaP spheroids for RNA extraction, 30 spheroids/BR of 3DS and 10 spheroids/BR of 3DF spheroids were collected in triplicate and washed twice with PBS +/+ (Gibco). For 2D LNCaP culture, samples were washed twice with PBS +/+. LNCaP samples were lyzed with the TRI Reagent (ZYMO Research). RNA extraction is then performed using the DirectzolTM RNA MiniPrep Plus kit (ZYMO Research). Quality and Concentration of the extracted RNA solution were assessed with Thermo Scientific™ NanoDrop™ 2000/2000c Spectrophotometers. Triplicate RNA samples from each condition were diluted to 20 ng/*μ*l with DNase/RNase-Free Water (ZYMO RESEARCH). 15*μ*l RNA solution of each replicate were sent to the UCLA Center for Systems Biomedicine for RNA expression assay with the Nanostring™ nCounter. Expression of 45 genes related to metabolic pathway, androgen receptor signaling, hypoxia, viability, are neuroendocrine differentiation were analyzed. The results were normalized to the average gene expression of three housekeeping genes. Principal Component Analysis and volcano plots were created in OriginPro for gene expression visualization.

### Transmitted Light Microscopy and Fluorescent Imaging

All stained LNCaP samples, shown in figures 1 and 3, were imaged using a Confocal microscope with a 10x or 40x objective. Cell viability images (figures 2 and 5) were acquired with an inverted microscope (Etaluma LS720) with a 4x phase contrast objective (Olympus, UPLFLN4XIPC) inside an incubator. For each well of the 96 wells, one image for each channel (i.e., phase contrast, 405 nm, and 488 nm) was obtained with a field of view ~ 770*μ*m×770*μ*m. To conduct the time-lapse experiment, every well was imaged every 20 minutes over a period of 48 hours. While our selected antibodies have been previously validated, we also examined the non-specific binding by measuring the fluorescent intensity in samples that were only stained with secondary antibodies. Prior to further analyses, the background of the fluorescent data was evaluated and subtracted. Image analysis was performed using Fiji Image J and Python.

### Immunoblot Analysis

PC3 and LNCaP cells were obtained from ATCC and cultured according to manufacturer’s protocols. LNCaP 2D cells were collected after 72 hours of treatment with DMSO or Enzalutamide (10 uM) and LNCaP aggregates were collected after 7 days in Static or Flow culture. All cells were lysed in RIPA buffer (50mM Tris-HCl pH 8.0, 150mM NaCl, 1% NP-40, 0.5% Sodium Deoxycholate, 0.1% SDS) containing a complete protease inhibitor cocktail tablet (Roche). Every sample was sonicated with sonic dismembrator (Fisher) to improve nuclear and membranous protein extraction. Proteins were run on NuPAGE 4%–12% Bis-Tris Gel (Novex) and transferred onto PVDF membranes (Millipore Sigma) and probed with antibodies. PSA, ACTIN, ECAD, and NCAD were detected via fluorescence using goat anti-rabbit IgG-Alexa Fluor 647 (Invitrogen) or goat anti-mouse IgG-Alexa Fluor 647 (Invitrogen) and all others using HRP-conjugated antibodies against rabbit or mouse (Invitrogen). The full list of immunoblot antibodies and dilutions may be found in SI Table S3. Quantification of blots was performed in Fiji Image J.

### Statistical Analysis

Data are reported as mean values ± standard deviation (SD). Statistical analysis was performed in Microsoft Excel using 2-tailed Student’s T-Tests. Significance levels are indicated with asterisks in each figure caption. P values ≤ 0.05, 0.01, and 0.001 are indicated with one, two, or three asterisks, respectively. PCA, volcano plots, and bar charts were performed in Origin Pro.

## Funding

This work was funded by the Broad Stem Cell Research Center at the University of California, Los Angeles, the UCLA SPORE in Prostate Cancer Grant (P50 CA092131), and the Prostate Cancer Foundation Young Investigator Award. A.S.G. is supported by the National Cancer Institute of the National Institutes of Health under Award Number R01CA237191, by University of California’s Cancer Research Coordinating Committee Award C23CR5598, by American Cancer Society award RSG-17-068-01-TBG, the UCLA Eli and Edythe Broad Center of Regenerative Medicine and Stem Cell Research Rose Hills Foundation Innovator Grant, the UCLA Jonsson Comprehensive Cancer Center and Eli and Edythe Broad Center of Regenerative Medicine and Stem Cell Research Ablon Scholars Program, the National Center for Advancing Translational Sciences UCLA CTSI Grant UL1TR001881, STOP CANCER, and the UCLA Institute of Urologic Oncology. This material is also based upon work supported by the National Science Foundation Graduate Research Fellowship under Grant No. DGE-2034835.

We are grateful to Dorian Luccioni and Ethan Salter for assistance in the engineering renderings.

## Author Contributions

M.-C.P., A.-S.G., and N.-Y.C.L. designed the project. M.-C.P., A.-S.G., and N.-Y.C. L. wrote the manuscript. M.-C.P., S.H., T.H., S.I., B.-S.L., M.-J.R., J.-A.D., and N.-Y.C. performed the experiments. M.-C.P., T.H., S.I., M.-J.R., J.-A.D., and N.-Y.C. analyzed the data. M.-J.R. performed RNA extraction & J.-A.D performed Western Blot. T.H. and A.-S.G. provided the LNCaP cells. All authors reviewed and approved the manuscript.

## Competing Interests

The authors declare no commercial or financial conflict of interest.

## Data and materials availability

All information needed to evaluate the conclusions in the paper are present in the paper and/or the Supplementary Materials. All data needed to reproduce the figures in the manuscript are available online at https://doi.org/10.5281/zenodo.7542326.

## Supplementary materials

Figures S1-S14

Tables S1-S3

Captions for Video S1 Supplementary Video S1

## Notes

https://doi.org/10.5281/zenodo.7542326

### Competing Interest Statement

The authors have declared no competing interest.

### Summary of Updates

Figures 1 and 5 revised; Additional data to strengthen comparisons between other flow devices and to validate necrotic phenotype in static samples; Supplemental files updated; authors updated.

