## Supplementary Information for "Microwell-Based Flow Culture Increases Viability and Restores Drug Response in Prostate Cancer Spheroids"

**This PDF file includes:**

Figs. S1-S14
Caption for Table S1
Tables S2-S3
Caption for Video S1

**Other Supplementary Materials for this manuscript include the following:**

Table S1
Video S1

**Figure S1.**

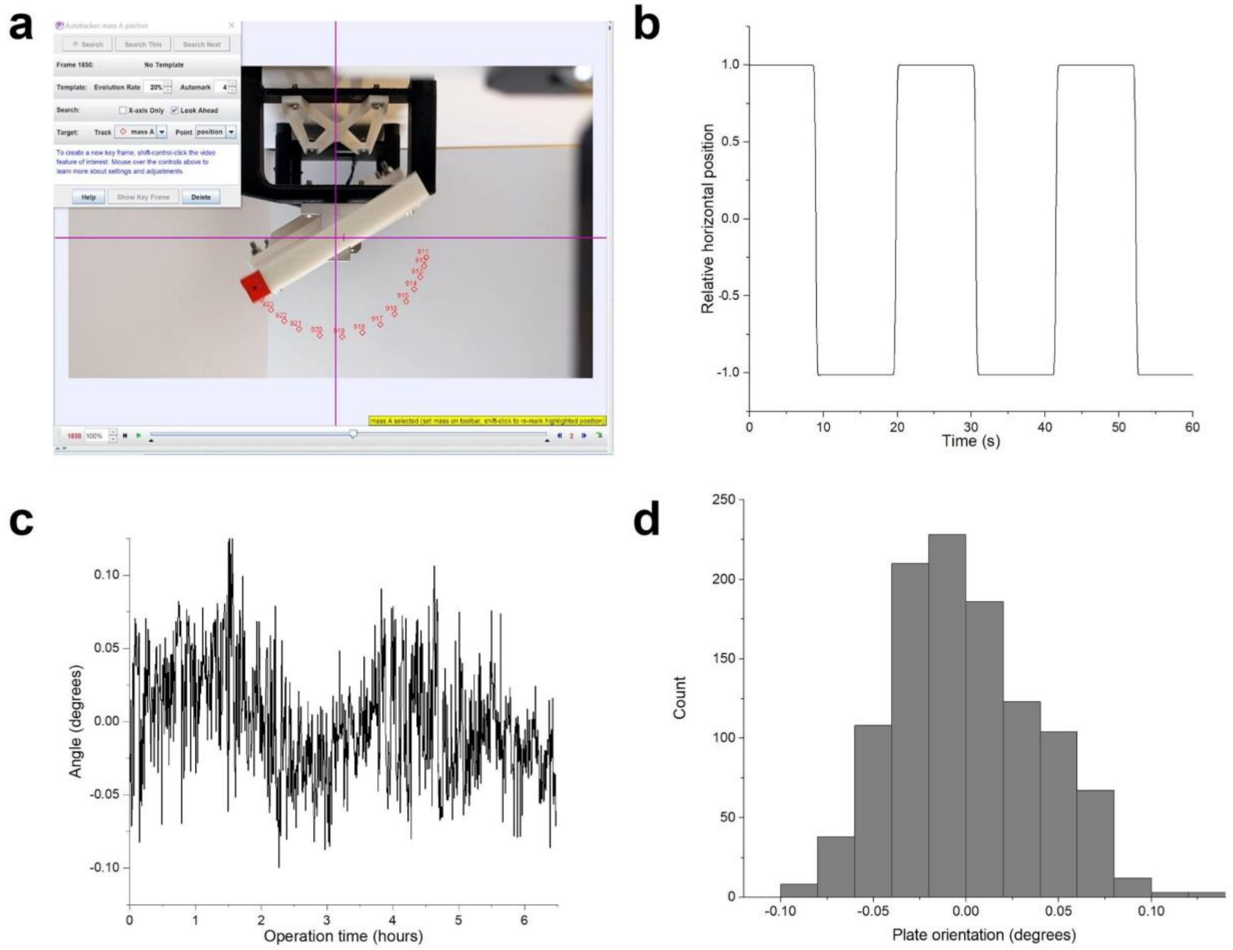


**Precision of rotation device. a** We characterized the precision of the rotation device through automated video analysis of a dot affixed to the outer edge of the rotating plate (Physlets Tracker). **b** The tracked marker stopped at the same horizontal position for each rotation. **c-d** Tracking the device's resting position over many cycles revealed no systematic deviation.

**Figure S2.**

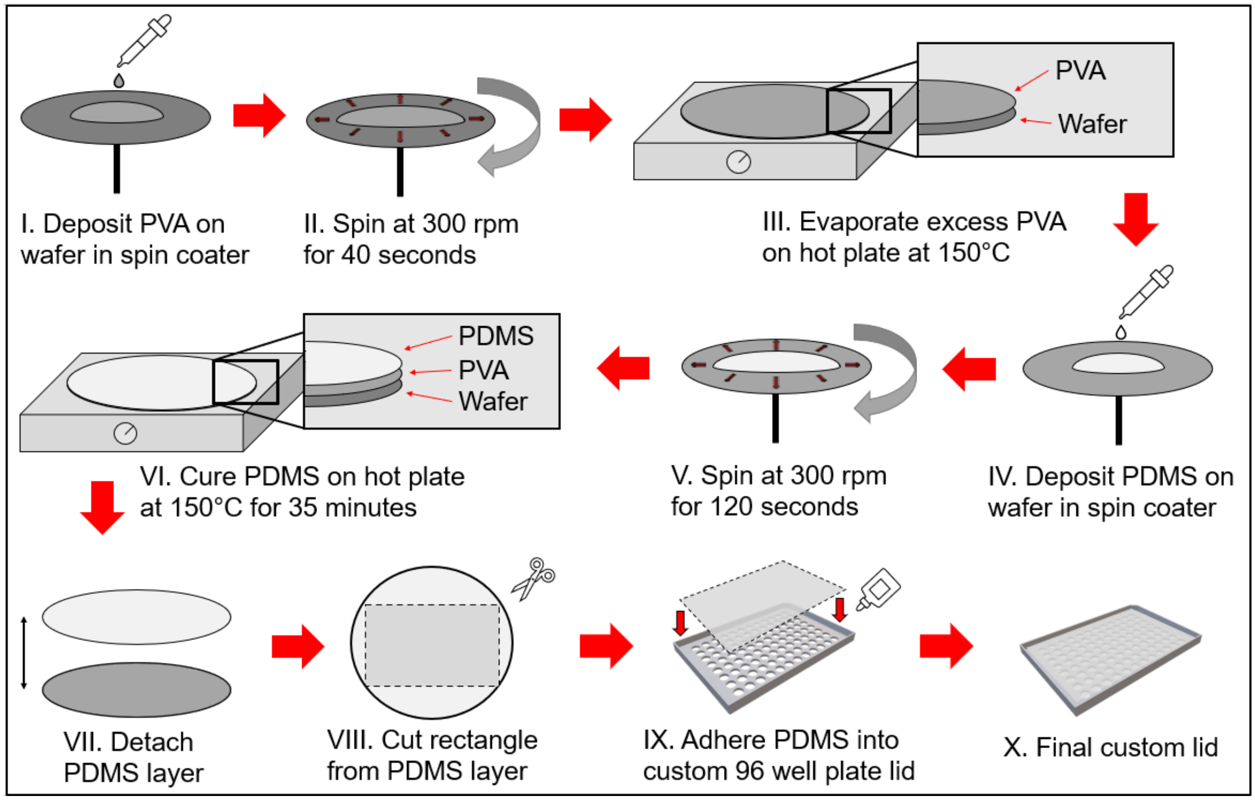


**Fabrication of PDMS membrane.** A sacrificial layer of polyvinyl alcohol (PVA) is deposited on a silicon wafer (I) and spun at 300 rpm for 40 seconds to create an even layer (II). Excess PVA is evaporated off on a hot plate at 150°C (III). PDMS is then deposited on the wafer (IV) and spun at 300 rpm for 120 seconds to create a membrane around 150 µm thick (V). The PDMS membrane is cured for 35 minutes at 150°C (VI), then detached from the wafer (VII). The detached membrane is cut to the shape of the custom lid (VIII) and adhered to the lid (IX) to create the final product (X).

**Figure S3.**

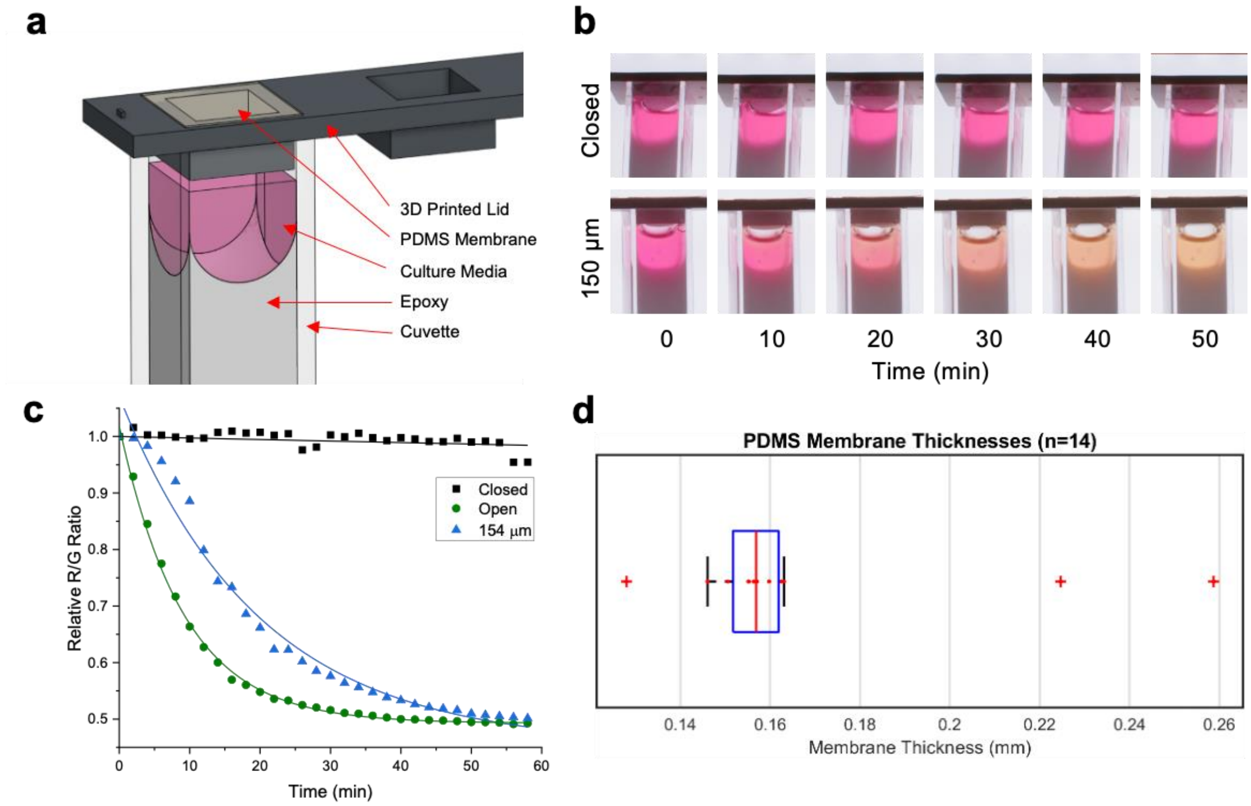


**PDMS membrane diffusivity comparison. a** Simulation of a 96-well plate well using an epoxy-filled cuvette to mimic the equivalent surface area-to-volume ratio. The opening at the surface allows for adhesion of a PDMS membrane for the experimental condition. **b** Closed, open, and experimental (150 $\mu m$, open image not shown) cuvettes were put into an incubator with a 40% $CO_2$ concentration and imaged over time. **c** The red/green color ratio of the pH indicator in the culture media was quantified as a proxy for $CO_2$ diffusion, showing that the PDMS had a CO2 diffusivity comparable to that of an open cuvette. **d** The fabrication of PDMS membranes at a specific thickness had a small error margin.

**Figure S4.**

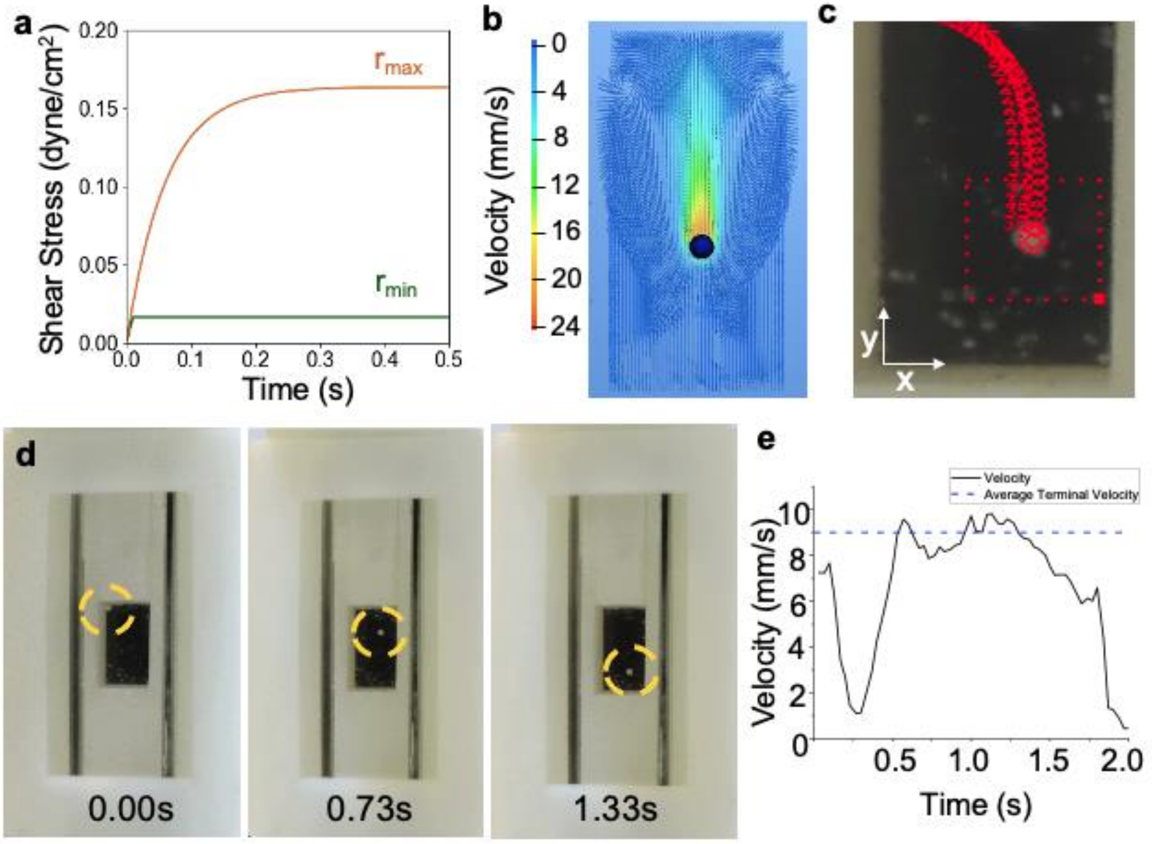


**Spheroid pathway tracking. a** Mathematical model of shear stress over time as the spheroid ($r_{min}$ = 50$\mu m$ and $r_{max}$ = 500$\mu m$) reaches terminal velocity. In the calculation we assumed a media density of 1000 kg/m$^3$, a viscosity of 1x10$^{-3}$ Pa-s, and an organoid density of 1100 kg/m$^3$. **b** COMSOL fluid simulation of an idealized spheroid (black sphere) during sedimentation in a well. Simulation arrows represent fluid displacement around the spheroid, which is at terminal velocity. **c** Spheroid (r = 500 µm) settling pathway with position tracking (Physlets Tracker). d Snapshots of spheroid at different timepoints in settling pathway. **e** Spheroid velocity over time for duration of motion. Dashed line marks the average terminal velocity, which corresponds to a shear stress of ~0.16 dynes/cm$^2$.

**Figure S5.**

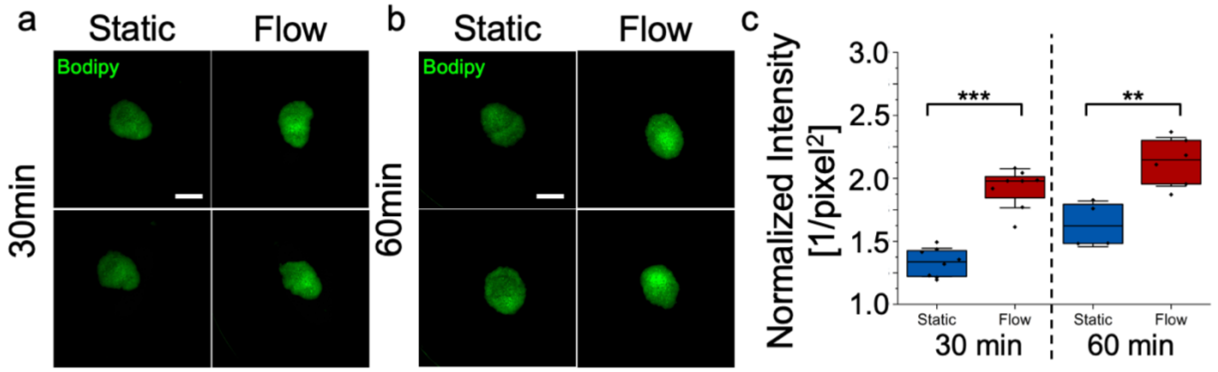


**Bodipy diffusion in LNCaP spheroids at 30min and 60min timepoints. a-b** Spheroids were aggregated for 3 days following the steps described in the main manuscript. Bodipy ($5\boldsymbol{\mu g/ml}$) was added to each sample, and the spheroids were cultured in flow or static conditions for 30 minutes (mins) (**a**) or 60 mins (**b**). **c** Following incubation, the samples were fluorescently imaged and the Bodipy intensity was quantified from the corresponding images (30 min: static and flow n = 2 biological replicates (BRs), 3 technical replicates (TRs)/BR; 60 min: static and flow n = 3 BRs, flow n = 2 TRs/BR and static n = 1-2 TRs/BR). At both time points, flow shows an increase in Bodipy signal (*** $p < 0.001$, ** $p < 0.01$, two-tailed Student's T-Test, scale bar = 500 $\boldsymbol{\mu m}$).

**Figure S6.**

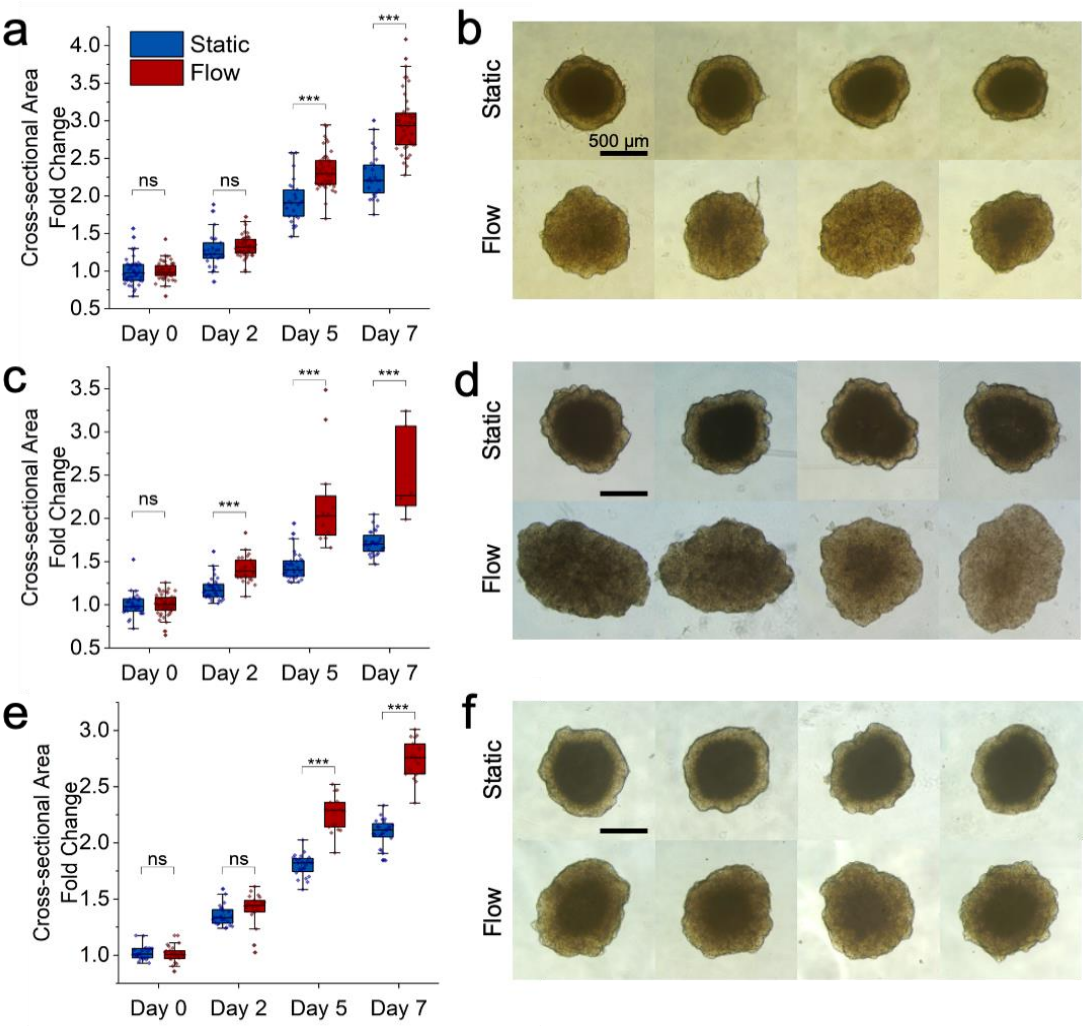


**Cross-sectional area replicates in flow and static culture.** This figure shows replicates for Figure 1f of the main manuscript. **a, c,** and **e** show boxplots analyzing the fold change of the cross-sectional area of 3 separate experiment runs between the two tested culture conditions (**a** static n=25-42, flow n=44-48 BRs, **c** static n=28-34, flow n=6-47 BRs, **e** static n=24 BRs, flow n=19-24 BRs). All brightfield images in **b, d,** and **f** show spheroids on day 7 of static or flow culture, respectively. (ns = non-significant; * $p < 0.05$; ** $p < 0.001$; *** $p < 0.0001$, two-tailed Student's T-Test).

**Figure S7.**

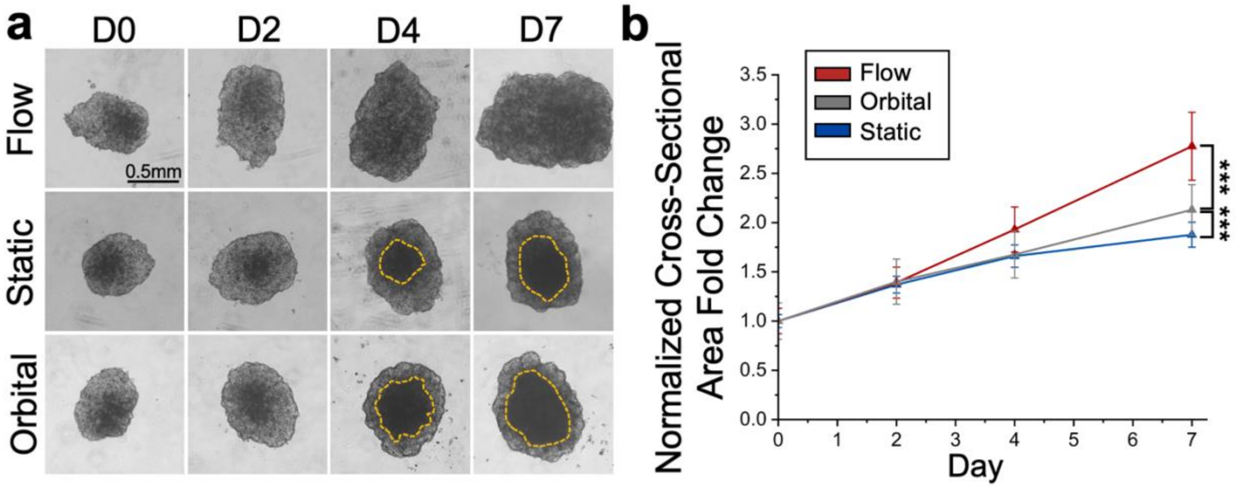


**Comparison of flow culture with orbital shaker and static culture. a** Brightfield images of MFD (flow), static, and orbital shaker (orbital) samples cultured over 7 days. 5k cells/well were seeded in 96 v-bottom plates and allowed to aggregate for 3 days at 37C. On day 0 (D0), samples were transferred into 96 u-bottom plates and separated into the MFD, static, and orbital shaking (100rpm) conditions. The necrotic core (absent in flow samples) on D4 and D7 is outlined in yellow. **b** Quantification of spheroid growth over 7 days. Each data point represents the average sample size normalized to the average of the respective condition on D0. Significance stars between sample averages on D7 indicate $p < 0.001$, two-tailed Student's T-Test. D7 static n=23, flow n=23, orbital n=13 BRs.

**Figure S8.**

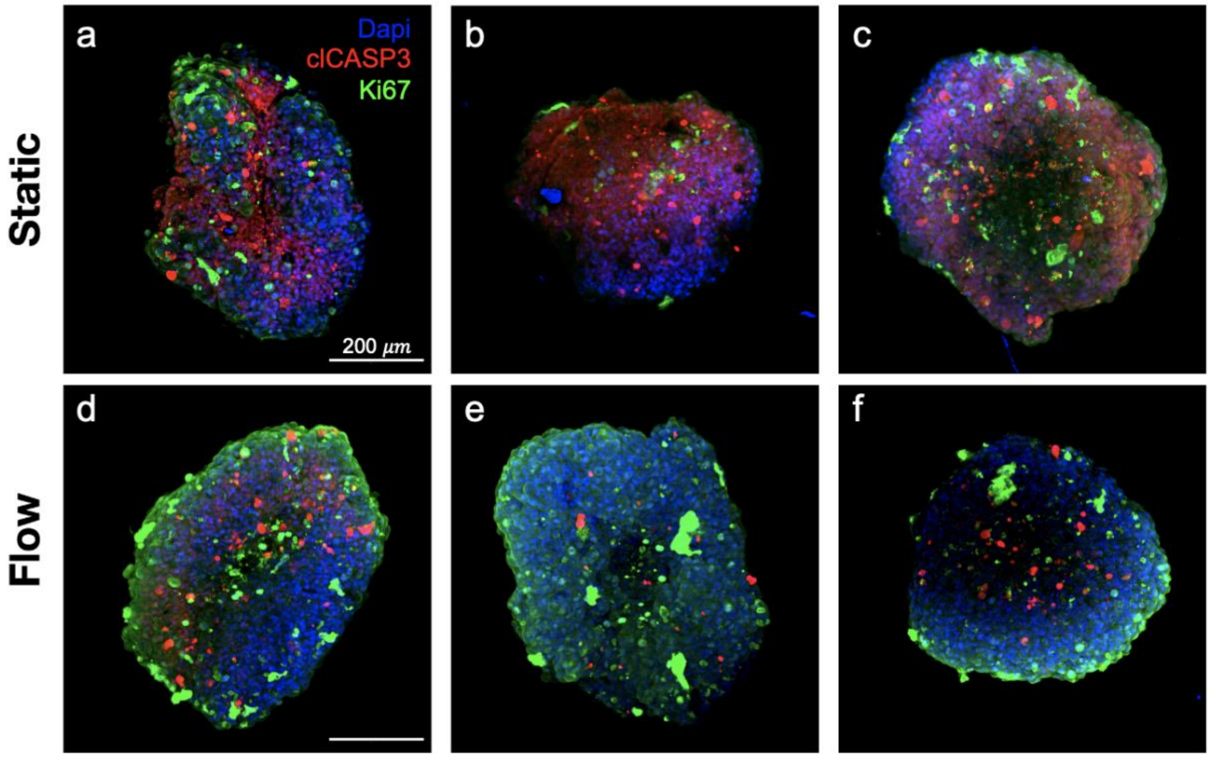


**Immunofluorescent replicates of cellular proliferation and apoptosis in flow and static samples.** This figure shows replicates for Figure 1h and 1i of the main manuscript. **a-f** 3DS samples display greater expression of clCASP3 (cell apoptosis) and reduced Ki67 (cell proliferation) compared to 3DF samples. (Flow and static n = 3 BRs).

**Figure S9.**

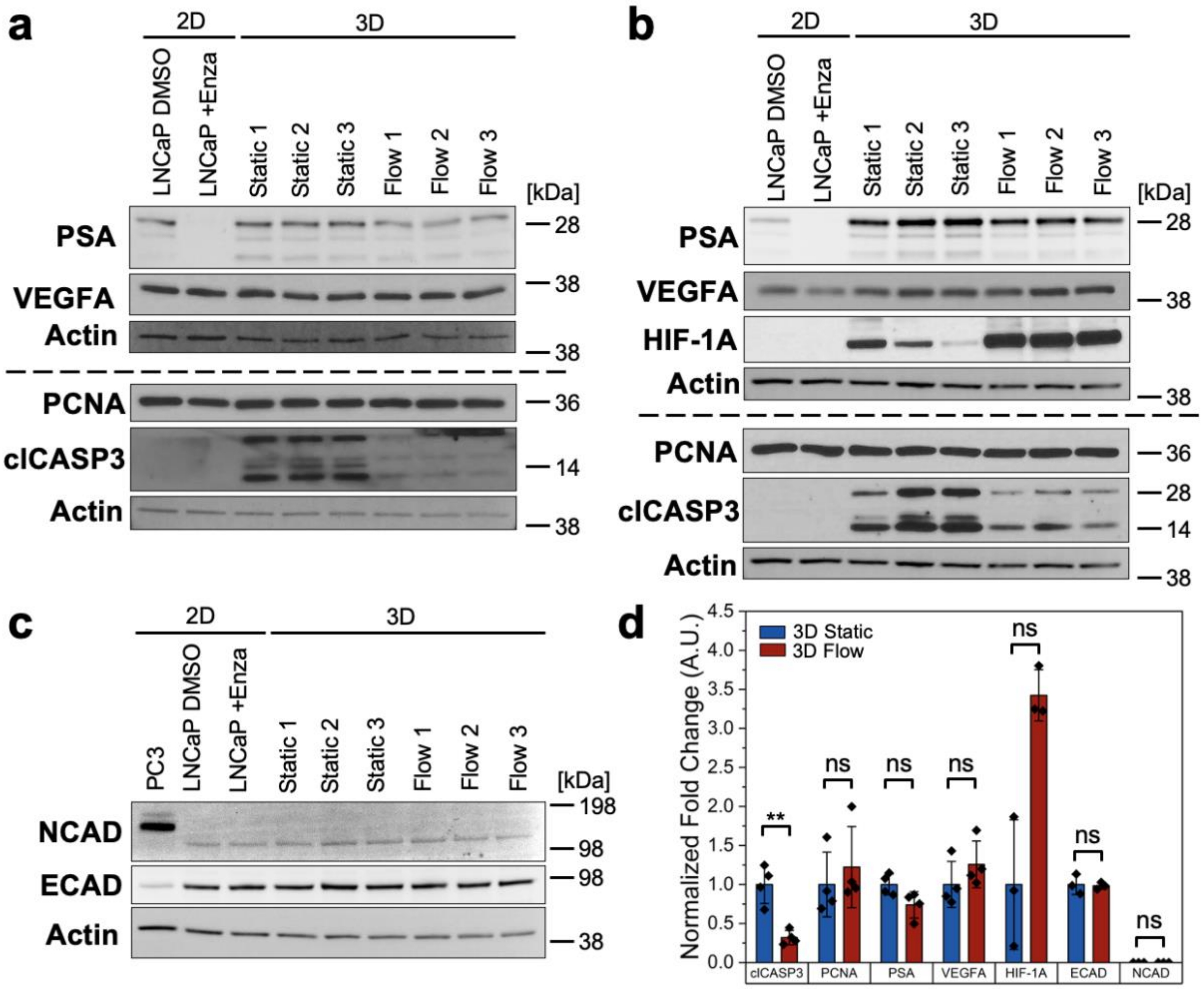


**Western Blot Analysis of PCa, hypoxia, and EMT protein expression. a-c** Western Blot analysis for D7 LNCaP spheroids in flow and static with positive and/or negative 2D controls. LNCaP + Enza refers to cells treated with androgen receptor inhibitor Enzalutamide. (**a** 3D static and flow 3 TRs; **b-c** 3D static & flow 3 BRs). **d** Quantification of relative 3D static and flow protein expression (Image J). All TRs shown in **a** were averaged to form one BR. (ns = non-significant; ** p<0.01, two-tailed Student's T-Test).

**Figure S10.**

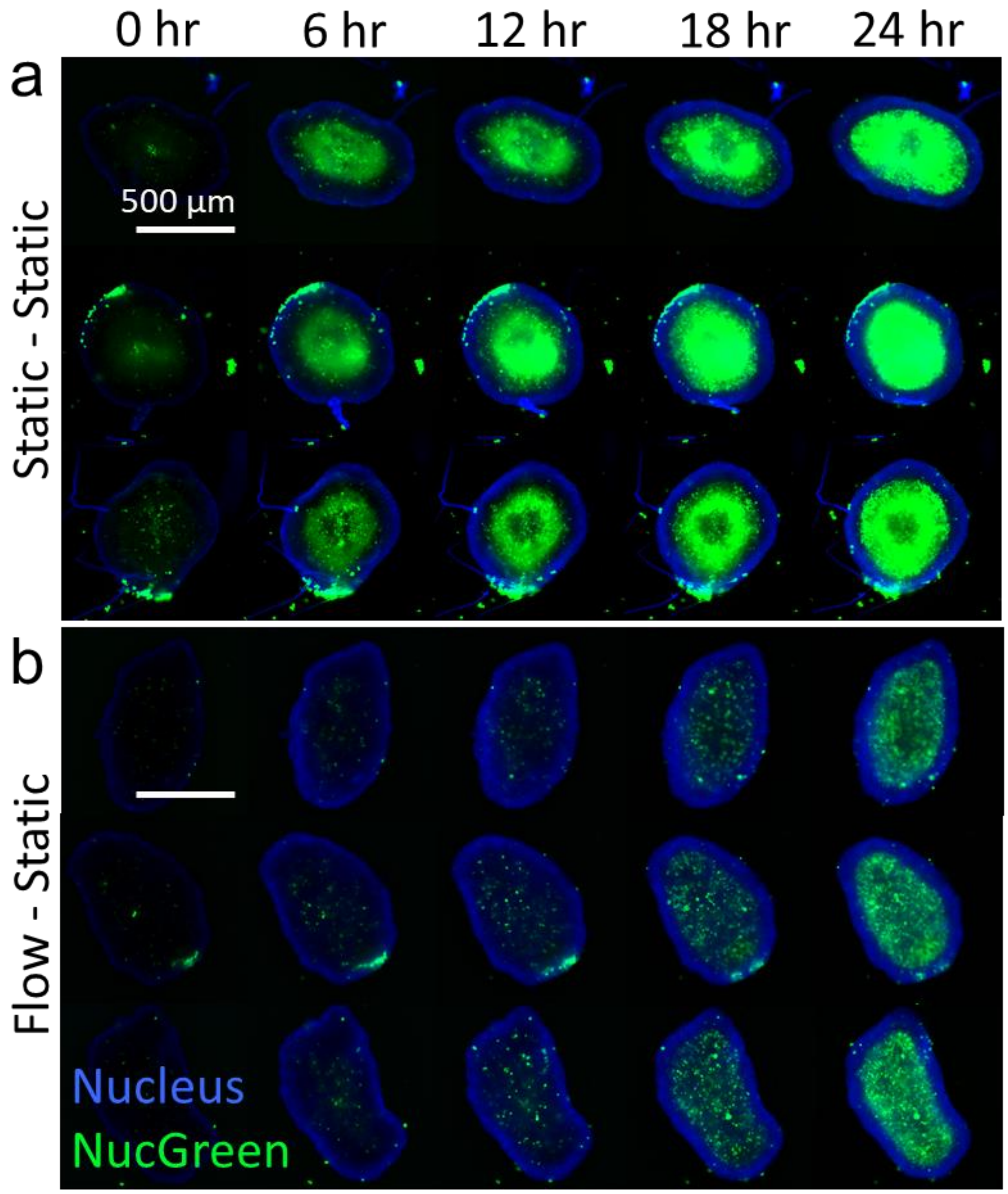


**24-hour untreated timelapse replicates for flow and static conditions.** This figure shows BRs for figure 2 in the main manuscript. Static (**a**) and flow (**b**) samples were placed in static culture on D7 for 24 hours. The samples were stained for total cells (NucBlue) and dead cells (NucGreen) at 0 hours and imaged every 20 minutes at 37C (Etaluma, 4x).

**Figure S11.**

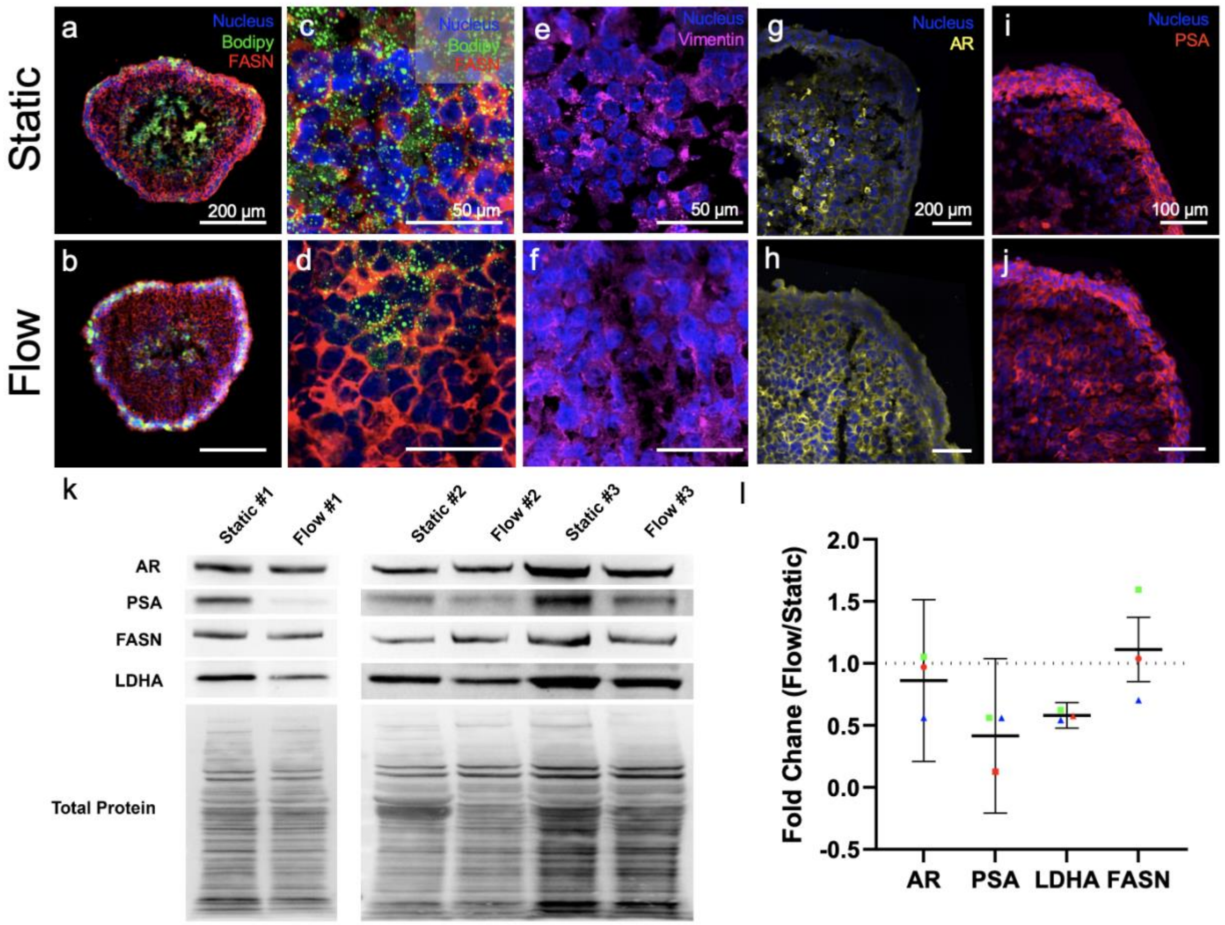


**Supplementary immunostaining and Western Blot for PCa-related protein markers. a-d** Immunostained LNCaP sections for Bodipy and FASN to visualize lipid droplets and fatty acid production, respectively. **e-f** Static and flow samples express Vimentin, an intermediate filament protein. **g-h** Immunofluorescent imaging of androgen receptor (AR). **i-j** Immunofluorescent imaging of prostate specific antigen (PSA). **k** Western blot (WB) results for AR, PSA, FASN, and lactate dehydrogenase A (LDHA). **l** Quantification of relative flow and static WB protein expression. All values were normalized to the total protein expression. The fold change of flow compared to static for three independent runs is plotted (n = 1 for each data point). All protein expression is non-significant between static and flow.

**Figure S12.**

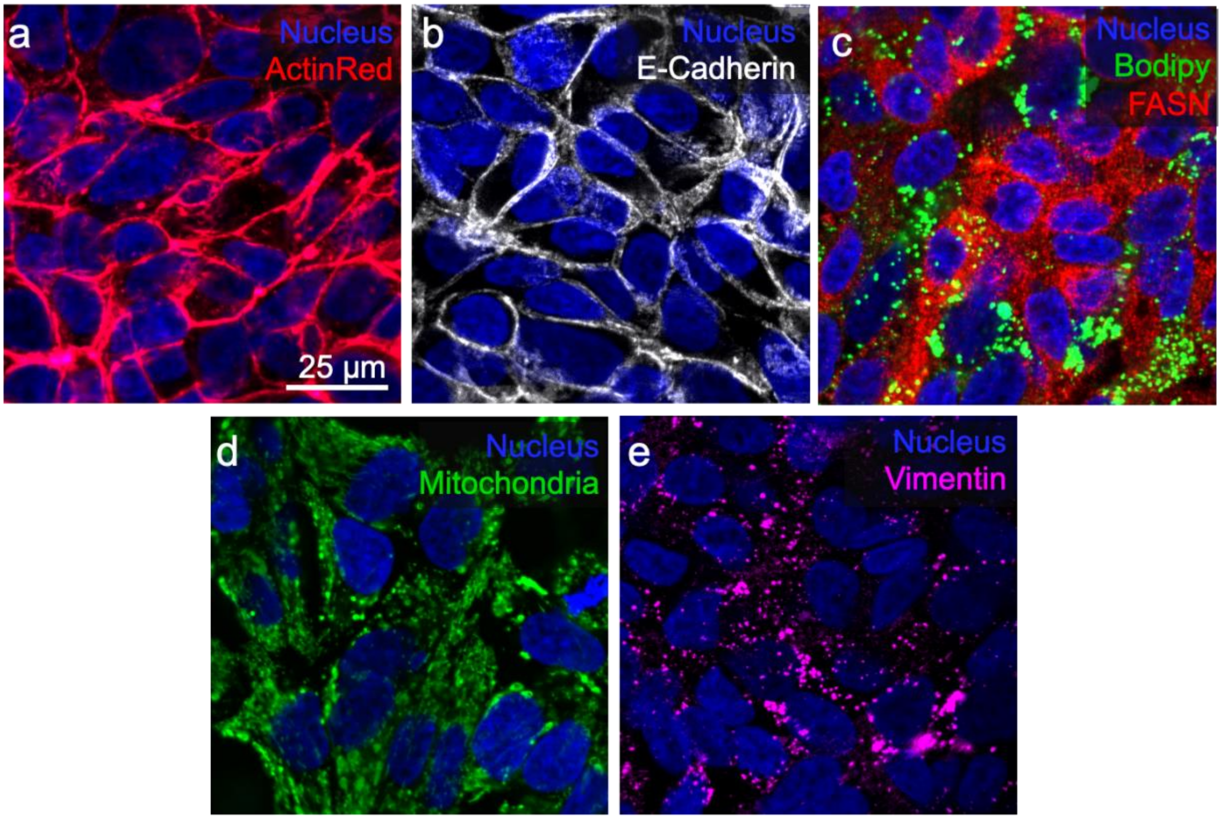


**Validation of staining reagents in 2D control culture. a-e** 2D LNCaP culture was stained for Actin, E-cadherin, Bodipy, FASN, Mitochondria, and Vimentin as positive controls for comparison with 3D sectioned LNCaP data.

**Figure S13.**

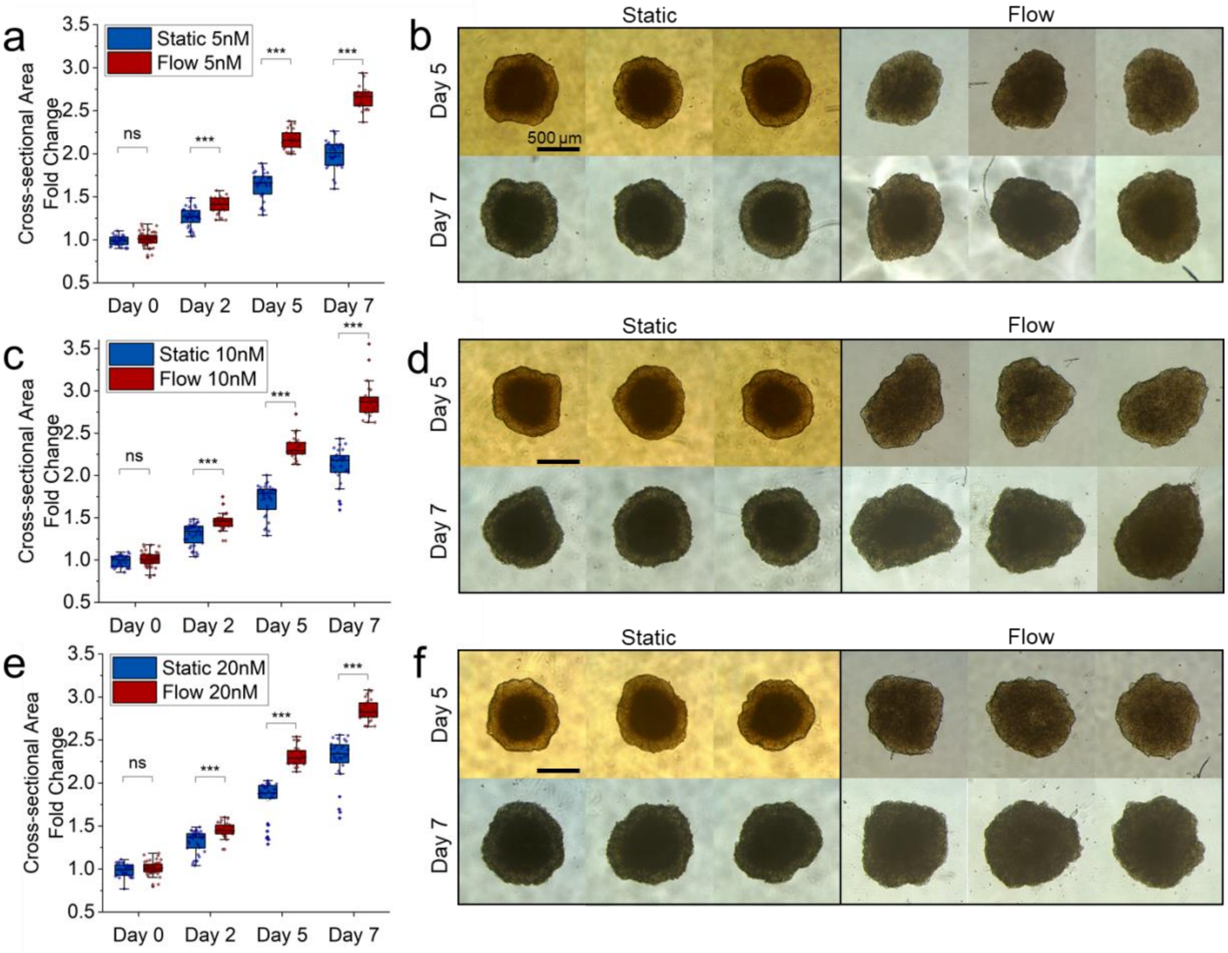


**Docetaxel treatment morphology images. a-f** Docetaxel (Selleck Chemicals, S1148) was administered at 0, 5, 10 and 20nMs for 48 hours from day 5 (**a** static n = 23-24, flow = 21-24 BRs; **c** static n = 24, flow = 24 BRs; **e** static n = 24, flow = 23-24 BRs; ns = non-significant; *** p < 0.001, two-tailed Student's T-Test).

**Figure S14.**

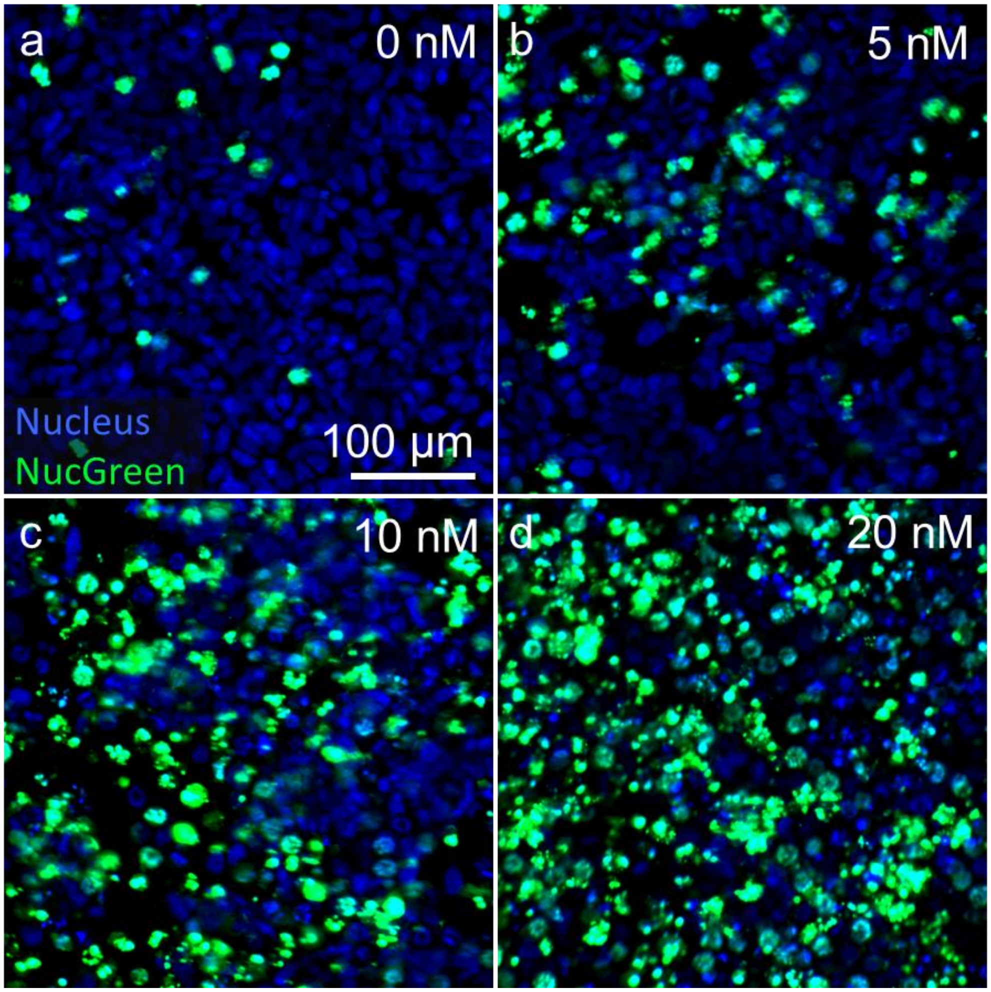


**Docetaxel 2D Control at varying dosages.** Staining for all cells (NucBlue – Nucleus) and dead cells (NucGreen) shows increasing cell death at higher concentrations of Docetaxel.

**Table S1.**

**Nanostring Gene Panel.** We identified 45 genes including 3 housekeeping genes that we hypothesized could be impacted by flow culture. This selection was based on previous studies which categorized cells/spheroids from prostate cancer cell lines based on common PCa markers or introduced metrics to analyze cellular response to metabolic or hypoxic stress. The selected genes comprised PCa, metabolic pathway, androgen receptor pathway, cell viability, hypoxia, and neuroendocrine markers.

**Table S2.**

a

| Antibody or Stain | Source | Catalog # | Host species, Isotype | Dilution |
|---|---|---|---|---|
| PSA | Cell signaling | 5877 | Rabbit, IgG | 1:400 |
| Vimentin | Invitrogen | MA5-11883 | Mouse, IgG1 | 1:250 |
| Androgen receptor | Invitrogen | MA5-13426 | Mouse, IgG1 | 1:40 |
| E-Cadherin | Invitrogen | 14-3249-82 | Rat, IgG1 | 1:100 |
| Cleaved Caspase-3 | Cell signaling | 9661 | Rabbit | 1:400 |
| Mitochondria | Invitrogen | MA5-12017 | Mouse, IgG1 | 1:200 |
| TUFM | Atlas Antibodies | AMAb90966 | Mouse, IgG1 | 1:200 |
| Ki67 | Invitrogen | MA5-15690 | Mouse, IgG1 | 1:400 |
| FASN | Bethyl Laboratories | 50-157-1866 | Rabbit | 1:300 |
| Bodipy | Invitrogen | D3922 | - | 5 µg/ml |
| ActinRed | Invitrogen | R37112 | - | 2 drops/ml |
| Hypoxia Tracker | Invitrogen | I14833 | - | 2.5 µM |
| ROS Tracker | Invitrogen | C10422 | - | 1:500 |
| NucGreen | Invitrogen | R37609 | - | 2 drops/ml |
| NucBlue | Invitrogen | R37606 | - | 1 drop/ml |

b

| Secondary Antibody | Source | Catalog # | Wavelength | Dilution |
|---|---|---|---|---|
| Donkey anti-Rabbit | Invitrogen | A32795 | 647 nm | 1:1000 |
| Donkey anti-Mouse | Invitrogen | A32766 | 488 nm | 1:1000 |
| Goat anti-Rat | Invitrogen | A21247 | 647 nm | 1:1000 |
| Donkey anti-Goat | Jackson ImmunoResearch | 705-165147 | Cy3 | 1:1000 |

**Immunostaining reagents. a** List of primary antibodies and stains used for immunofluorescent imaging with the respective dilution ratio. **b** Table summarizing information about the secondary antibodies used.

**Table S3.**

| Antibody | Source | Catalog # | Host Species, Isotype | Dilution |
|---|---|---|---|---|
| AR | Cell Signaling | 5153 | Rabbit | 1:2000 |
| PSA | Cell Signaling | 5877 | Rabbit | 1:1000 |
| PCNA | Cell Signaling | 2586 | Mouse | 1:2000 |
| clCASP3 | Cell Signaling | 9661 | Rabbit | 1:250 |
| VEGFA | Abcam | ab46154 | Rabbit | 1:1000 |
| HIF-1$\alpha$ | Cell Signaling | 14179 | Rabbit | 1:1000 |
| ECAD | ProteinTech | 20874-1-AP | Rabbit | 1:1000 |
| NCAD | ThermoFisher | 33-3900 | Mouse | 1:1000 |
| ACTIN | ThermoFisher | PA1-16889 | Rabbit | 1:20000 |
| ACTIN | ThermoFisher | PMIA1140 | Mouse | 1:20000 |
| **Secondary Antibody** | **Source** | **Catalog #** | **Wavelength** | **Dilution** |
| Goat anti-Rabbit, IgG | ThermoFisher | A21244 | 647 nm | 1:1000 |
| Goat anti-Mouse, IgG | ThermoFisher | A21235 | 647 nm | 1:1000 |
| Goat anti-Rabbit, IgG (H+L) HRP | ThermoFisher | 31463 | - | 1:10000 |
| Goat anti-Mouse, IgG (H+L) HRP | ThermoFisher | 31430 | - | 1:10000 |

**Immunoblot Antibodies.** Table of all primary and secondary antibodies used for Western Blot analysis.

**Video S1.**

**Necrotic core formation after flow cessation.** This video corresponds to Figure 3 in the main manuscript and Supplemental Figure 7. Flow and static LNCaP samples were placed in static culture and live stained with NucGreen (dead cells) and NucBlue (cell nucleus). The flow-static and static-static samples were time-lapsed imaged in an incubator every 20 minutes for 24 hours. A necrotic core emerges in the flow samples at 18 hours after stopping flow.
